# Discovering precise temporal patterns in large-scale neural recordings through robust and interpretable time warping

**DOI:** 10.1101/661165

**Authors:** Alex H. Williams, Ben Poole, Niru Maheswaranathan, Ashesh K. Dhawale, Tucker Fisher, Christopher D. Wilson, David H. Brann, Eric Trautmann, Stephen Ryu, Roman Shusterman, Dmitry Rinberg, Bence P. Ölveczky, Krishna V. Shenoy, Surya Ganguli

## Abstract

Though the temporal precision of neural computation has been studied intensively, a data-driven determination of this precision remains a fundamental challenge. Reproducible spike time patterns may be obscured on single trials by uncontrolled temporal variability in behavior and cognition, or may not even be time locked to measurable signatures in either behavior or local field potentials (LFP). To overcome these challenges, we describe a general-purpose time warping framework that reveals precise spike-time patterns in an unsupervised manner, even when spiking is decoupled from behavior or is temporally stretched across single trials. We demonstrate this method across diverse systems: cued reaching in nonhuman primates, motor sequence production in rats, and olfaction in mice. This approach flexibly uncovers diverse dynamical firing patterns, including pulsatile responses to behavioral events, LFP-aligned oscillatory spiking, and even unanticipated patterns, like 7 Hz oscillations in rat motor cortex that are not time-locked to measured behaviors or LFP.

## Introduction

The role of spike time precision in neural computation has been widely examined from both experimental and theoretical al perspectives (Softky and Koch 1993; London et al. 2010; Bruno 2011; Amarasingham et al. 2012; Amarasingham et al. 2015; Brette 2015; Denéve and Machens 2016), engendering intense debates in systems neuroscience over the last several decades. Empirically determining the degree of temporal precision from data is challenging because multi-neuronal spike trains may contain highly structured temporal patterns that are completely masked by temporal variations in behavioral and cognitive variables not under direct experimental control. For example, precise spike patterns may not be temporally locked to naïvely chosen sensory or behavioral events. Indeed, the fidelity of olfactory coding may be underestimated by factors of two to four when spike times are aligned to stimulus delivery instead of inhalation onset (Shusterman et al. 2011; Cury and Uchida 2010; Shusterman et al. 2018).

Thus, experimental estimates of spike time precision hinge on the choice of an alignment point, which defines the origin of the time axis on each trial. This choice can often be challenging and subjective. Even in relatively simple behavioral tasks, animals can experience a sequence of stimuli, actions, and rewards, each of which occur with varying latencies on different trials. Such tasks thus provide multiple choices for aligning multineuronal spike trains to measurable events marking an origin of time. Moreover, in addition to choosing an origin of time, we must also choose its units. Should spike times be measured in absolute clock time relative to some measured event, or in units of fractional time between two events? Should the units of time change between successive pairs of events? Could any one of these choices unmask spike-timing precision that is otherwise invisible?

Past studies have addressed these challenges in a number of ways: grouping trials together with similar durations before averaging spike counts (Murakami et al. 2014; Starkweather et al. 2017; Wang et al. 2018), manually stretching or compressing time units between measured task events (Leonardo and Fee 2005; Shusterman et al. 2011; Kobak et al. 2016; Aronov et al. 2017), or repeating statistical analyses around different choices of alignment point (Feierstein et al. 2006; Harvey et al. 2012; Jazayeri and Shadlen 2015; Shushruth et al. 2018). Determining an appropriate alignment and scaling of time is most challenging in systems far from the sensory or motor periphery, where neural responses are often not locked to any external event, and instead reflect internal decisions or changes-of-mind that occur at variable times within each trial. In these cases, the ideal temporal alignment point (e.g. the time a decision is made) may be entirely unmeasurable or ill-defined from the standpoint of behavior.

These complications, and the diversity of heuristic approaches used to address them, underscore a broad need for statistical frameworks to assess the temporal precision of neural computation. Of particular interest are *unsupervised* statistical methods that reveal precise patterns in multi-neuronal spike trains *without* reference to behavioral measurements. Such methods would be broadly applicable, as they make few assumptions about experimental design, animal model, or measured behaviors. Furthermore, by only considering the neural data, these methods may discover novel spike patterns aligned to unexpected variables. Most intriguingly, these methods could potentially identify spike patterns that are not well-aligned to any behavior, but instead to unobservable cognitive states, such as decision times.

While time series and image alignment methods are a well-studied topic in signal processing (Berndt and Clifford 1994; Marron et al. 2015; Mueen and Keogh 2016; Pnevmatikakis and Giovannucci 2017), these techniques have rarely been applied to large-scale neural recordings (but see recent work by Poole et al. 2017; Lawlor et al. 2018; Duncker and Sahani 2018). Neuroscientists have historically utilized simple alignment operations—namely, translating (Baker and Gerstein 2001; Ventura 2004) and potentially stretching/compressing activity traces between pairs of behavioral events (Shusterman et al. 2011; Leonardo and Fee 2005; Perez et al. 2013; Kobak et al. 2016). In contrast, popular statistical methods, such as *Dynamic Time Warping* (DTW; Berndt and Clifford 1994), allow signals to be non-uniformly compressed and dilated on each trial. While such *nonlinear warping* models can be useful, we demonstrate that they can be difficult to interpret and sensitive to the high level of noise that is typical of neural data.

To identify interpretable alignments for high-dimensional spike trains, we developed a framework for *linear* and *piecewise linear* time warping that encompass existing human-annotated procedures (Leonardo and Fee 2005; Kobak et al. 2016). Relative to nonlinear warping methods, the methods we describe are simpler to interpret, more robust to overfitting, and more computationally scalable. We applied these methods to multielectrode recordings collected from three experiments spanning different animal models (rodents and primates), brain regions (olfactory and motor cortex), and behavioral tasks (sensation and motor production). In each case, time warping revealed precise spike patterns that were imperceptible in the raw data. Moreover, some of these results were *not* easily captured by commonly chosen temporal alignments. For example, in rodents performing a motor timing task, we uncovered prominent *∼*7 Hz oscillations in spike times that were *not* aligned to the LFP or any measured behavioral event, providing a convincing example in which salient population dynamics would be likely overlooked by pre-defined behavioral alignments.

Overall, we demonstrate that simple time warping models can detect salient, yet easy-to-overlook, features in large-scale neural data. We expect these methods to particularly enable inquiry into circuits far from the sensory or motor periphery that are not tightly time-locked to any measurable stimulus or behavior. Yet, even in cases with seemingly obvious sensory or behavioral alignment points, our immediate findings suggest that common manual alignment and trial-averaging procedures may underestimate temporal precision, and may even qualitatively misrepresent single-trial dynamics. Thus, we provide a broadly applicable, data-driven framework to reveal and scientifically characterize fine-scale temporal features of neural dynamics. Such a framework, when combined with future experiments, may help in adjudicating long-standing debates surrounding the temporal precision of neural computation.

## Results

### Time Warping Framework

Our ultimate goal is to identify dynamical firing patterns that reliably occur on a trial-by-trial basis. If these activity patterns are tightly time-locked to a sensory or behavioral event, which can be confidently identified and measured, then we can accurately characterize the neural response by averaging over trials. This is illustrated in **Figure 1a** (left) which shows 100 trials of a simulated neural activity trace with additive Gaussian noise. The average activity trace (red trace; bottom) extracts the reliable features of the neural response from noisy single-trial instantiations (semi-transparent black traces). This synthetic example loosely resembles calcium fluorescence traces, but the methods we describe can be flexibly applied to any multi-dimensional time series including spike trains, fMRI data, or LFP traces.

More formally, if *N* neurons are measured at *T* timepoints over *K* nominally identical trials, the trial-average is given by: Here,

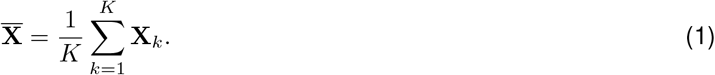

**Fig 1.**
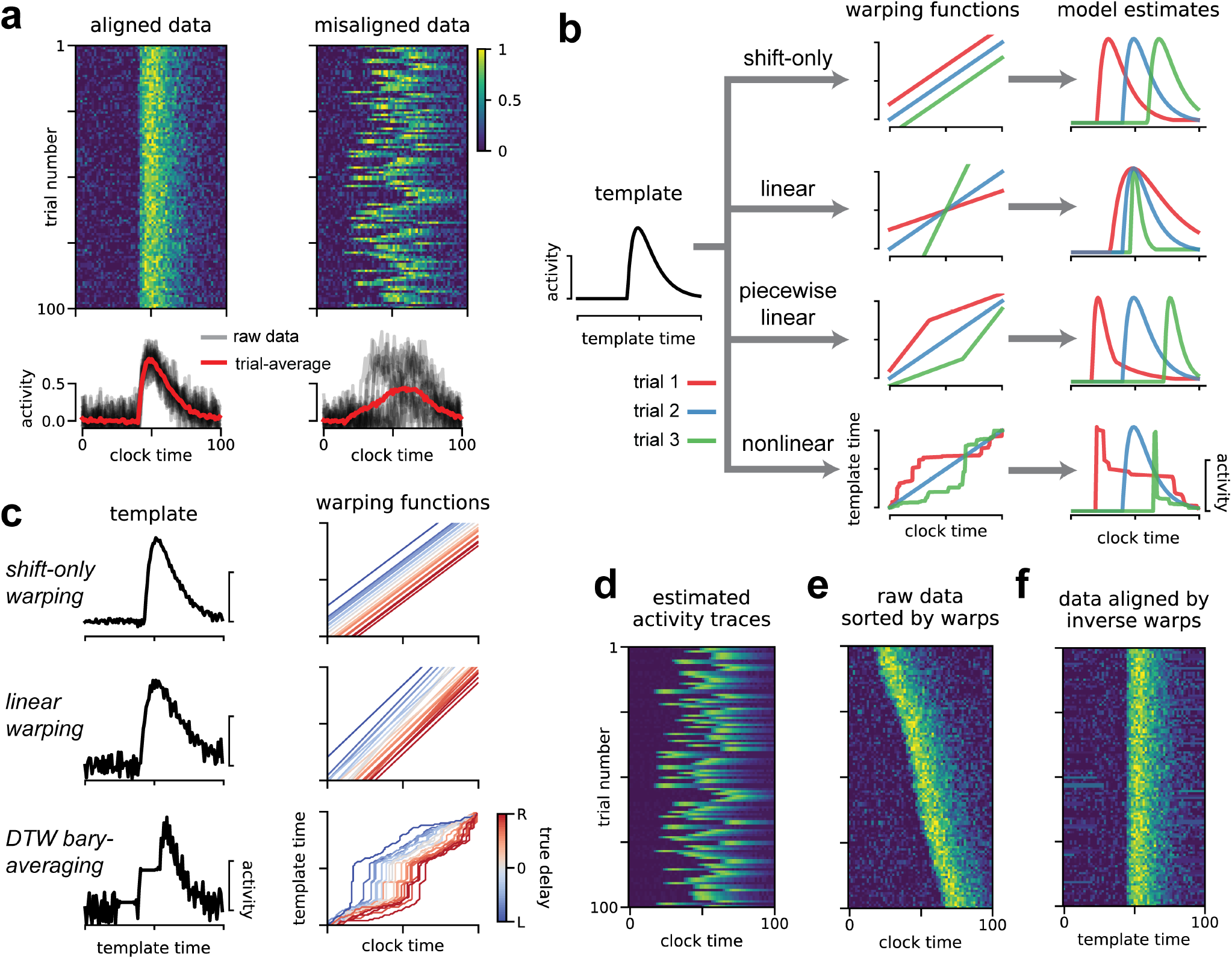
Illustration of time warping models. (A) Synthetic data from a single neuron on 100 trials. When the data are aligned (top left) the trial-average provides an effective, denoised description of activity (bottom left). When the data are misaligned by introducing jitter (right), the trial-average does not capture the typical firing pattern (bottom right). (B) Time warping models estimate a template (black line, left) that can be transformed on a trial-by-trial basis by warping functions (middle column) producing single-trial estimates of neural activity (right column). Red, blue and green lines represent warping functions and estimated firing rates on three example trials. Top row illustrates shifting the template activity in time (shift-only warping), second row illustrates stretching and compressing the template (linear warping), third row illustrates piecewise-linear warping with two line segments (one may increase the number of segments to introduce more nonlinearity into the warping function), the bottom row illustrates a fully nonlinear warping. (C) Results of a shift-only warping model (top), a linear warping model (middle), and a nonlinear warping model (bottom; Petitjean et al. 2011) fit to the synthetic data from panel *A*. For each model, we show the warping templates (left, black lines) and warping functions (right, lines colored by ground truth shift to the right vs left). (D) The predicted firing rates of the shift-only warping model provide a denoised trial-by-trial estimate of neural activity. (E) The raw data sorted by the per-trial shift parameter learned by the shift-only warping model. (F) Data aligned by the shift-only model; the per-trial shift learned on the model template is applied in the opposite direction to the raw data.

Here, X_*k*_ is an *N* × *T* matrix denoting the measured activity of all neurons on trial *k*. In the context of spiking data, each row of 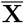 corresponds to a *peri-stimulus time histogram* (PSTH) of a recorded neuron. Trial averaging is also a common step in population-level statistical analyses (Gao and Ganguli 2015; Kobak et al. 2016).

Despite its widespread use, trial averaging can produce highly inaccurate and misleading results when neural activity is misaligned across trials. For example, introducing a random temporal shift to each simulated trial results in a less informative trial-average trace (**Fig 1a**, right). Such jitter commonly arises in practical applications, leading many research groups to develop custom-built alignment procedures for their experimental system. For example, in songbirds it is common to manually segment and cluster song syllables and warp intervening spike times on a per-syllable basis (Leonardo and Fee 2005). In olfaction, detailed measurements and fluid dynamics modeling of the sniff cycle have been pursued to understand the accuracy of sensory responses (Shusterman et al. 2011; Shusterman et al. 2018). Jitter and other forms of temporal variability are likely exacerbated in deeper brain areas, where there is greater opportunity for unobserved cognitive states (e.g. attentiveness) and actions (e.g. internal decision times) to influence the timing of dynamics.

Time warping methods address these challenges through a data-driven, statistical approach. The key idea is to fit a *response template* time series that is shifted and stretched—i.e., *warped*—on a trial-by-trial basis to match the data. The response template, denoted 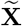, is an *N × T* matrix of activity traces that captures the average activity across trials *after* correcting for temporal deformations.

The time axis of the response template is then transformed by a *warping function* on each trial. Formally, we denote the warping function for trial *k* as *ω*_*k*_ (*t*); this function maps each timebin *t* (clock time) to a new index *ω*_*k*_ (*t*) (template time). If *ω*_*k*_(*t*) is an integer between 1 and *T*, then the warping transformation for every neuron *n* on trial *k* amounts to the transformation 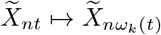 is *not* an integer then time warping is implemented by linear interpolation (see *Methods*). Note that this model assumes all recorded neurons share the same warping function on a trial-by-trial basis, though this could be relaxed by future work.

**Figure 1b** illustrates how different classes of warping functions account for single-trial variability in timing. In this paper, we focus on three main model classes: *shift-only time warping*, *linear time warping*, and *piecewise linear time warping* (**Fig 1b**, top three models). Shift-only warping represents the simplest model: the warping functions are constrained to be linear with slope equal to one, and only a single parameter (the y-intercept of *ω*_*k*_(*t*)) is fit on each trial. As its name suggests, the shift-only model can only account for trial-to-trial differences in response latency. In contrast, a linear warping model, which fits the slope in addition to the intercept of *ω*_*k*_(*t*), can account for variable latencies as well as uniform stretches or compressions of the response template. A piecewise linear warping model adds further complexity by adding one or more *knots* (points where the slope of *ω*_*k*_(*t*) can change). Most generally, nonlinear warping functions may be used, which non-uniformly stretch and compress portions of the template on each trial (**Fig 1b**, bottom).

In all cases, we constrain the time warping functions to be monotonically increasing. Intuitively, this ensures that the model cannot go backwards in time while making a prediction—that is, as *t* (clock time) increases, *ω*_*k*_ (*t*) (template time) must also increase. This ensures that the warping functions are invertible, which we later exploit to align data across trials. DTW-based time warping paths are not invertible, since the first derivative can be zero or infinite. Some other nonlinear warping methods (e.g. Duncker and Sahani 2018) do not require warping functions to be monotonic (and therefore invertible), though future work on these methods could incorporate this constraint.

The warping functions and response template are numerically optimized to minimize the total reconstruction error over all neurons, trials, and timepoints. For flexibility and computational efficiency, we chose the mean squared error to quantify reconstruction accuracy. Ignoring the interpolation step of time warping for the sake of clarity (see *Methods*), the total model error is:

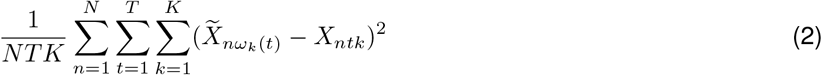

This expression is minimized with respect to the warping functions on each trial, *ω*_*k*_(*t*), and the response template, 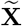. Other loss functions may be substituted for the mean squared error. In particular, a loss function based on Poisson noise is popular in neural modeling (Paninski 2004) and our accompanying Python package supports this option. To optimize the response template we utilize standard methods—least-squares solvers when minimizing squared loss, and gradient-based optimization when minimizing the Poisson objective function. To optimize the warping functions, we found that gradient-based methods were prone to converge to suboptimal local minima, and that it was preferable and tractable to use randomized searches over these low-dimensional parameter spaces (see *Methods*).

For illustration, we fit shift-only, linear, and nonlinear warping models to the misaligned synthetic data shown **Figure 1a**. By design, the shift-only model is sufficient to capture the ground truth variability in timing; as expected, this model identifies a highly accurate template firing pattern (**Fig 1c**, top left), along with warping functions that tightly correlate with the ground truth delay on a trial-by-trial basis (**Fig 1c**, top right). The linear warping model is a gentle extension of the shift-only warping model, which only introduces one additional parameter on each trial—the slope of each warping function. Yet, even this very minor extension produces a slightly worse estimate of the template and ground truth warping functions (**Fig 1c**, middle). This worsened estimate results from the model overfitting to noise in the simulated data; the linear warping model can use its additional per-trial parameter to align patterns of noise in the data across trials, which then appear in the response template. To demonstrate a more severe case of overfitting, we fit a nonlinear warping model using Dynamic Time Warping (DTW; Berndt and Clifford 1994), combined with a standard barycenter averaging procedure (Petitjean et al. 2011). This method can be highly effective on datasets with low levels of noise and complex temporal deformations. However, as we will soon see, neural datasets often exhibit the opposite—high levels of noise and simple temporal deformations. In this regime, DTW barycenter averaging identifies a noisy and deformed template (**Fig 1c**, bottom left) and warping paths that correlate with the ground truth jitter, but are unnecessarily complex (**Fig 1c**, bottom right).

These results demonstrate that time warping models can be sensitive to noise, especially when a more flexible class of warping functions is utilized. Our time warping framework uses three strategies to prevent overfitting. First, as illustrated by the progression of models in **Figure 1c**, we always compare the estimates of complex warping models (e.g. with piecewise linear warping functions) to the performance of simpler models (e.g. shift-only warping). Second, we include a smoothness regularization term on the template, which penalizes the average norm of the second temporal derivative, and thus encourages temporally smooth model estimates. Third, we place a penalty on the area between each warping function and the identity line, which penalizes the magnitude of warping on each trial. We include these roughness and warp-magnitude penalties in subsequent results, but show the results of unregularized time warping in **Figure 1c** for the sake of illustration. A detailed description of these regularization terms is provided in the *Methods* section.

These time warping models enable several strategies for visualizing and understanding neural data. First, one can directly inspect the model parameters (**Fig 1c**). The response template for each neuron captures the shape of the neural response, while the warping functions capture trial-to-trial variability. When simple warping functions are used, the parameters of each function (e.g. the slope and intercept, for a linear warping model) can be visualized in a histogram or scatterplot, or regressed against behavioral covariates. Second, one can view the model prediction as a denoised estimate of firing rates on a single-trial basis (**Fig 1d**). Third, one can re-sort the trials by the slope or the intercept of the warping function, producing a multi-trial raster plot that is easier to visually digest (**Fig 1e**). Finally, one can invert the learned warping functions on each trial to transform the raw data into an aligned time domain (**Fig 1f**). This alignment procedure simply entails plotting each activity trace as a function of 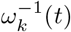 instead of raw clock time, *t*. Intuitively, this amounts to reversing the flow diagram shown **Figure 1b**, which is possible as long as the warping functions are monotonically increasing (i.e. invertible).

### Extraction of precise, ground truth spike patterns on synthetic data

Before proceeding to biological data, we examined a more challenging synthetic dataset involving multiple neurons and complex single-trial variability in timing. We simulated spike train data from *N* = 5 neurons, *T* = 150 timebins, and *K* = 75 trials. On each trial, the neural firing rates were time warped by randomized piecewise linear functions with one knot (the “ground truth” model; see *Methods*). This resulted in spike trains that appear highly variable in their raw form (**Fig 2a**).

**Fig 2.**
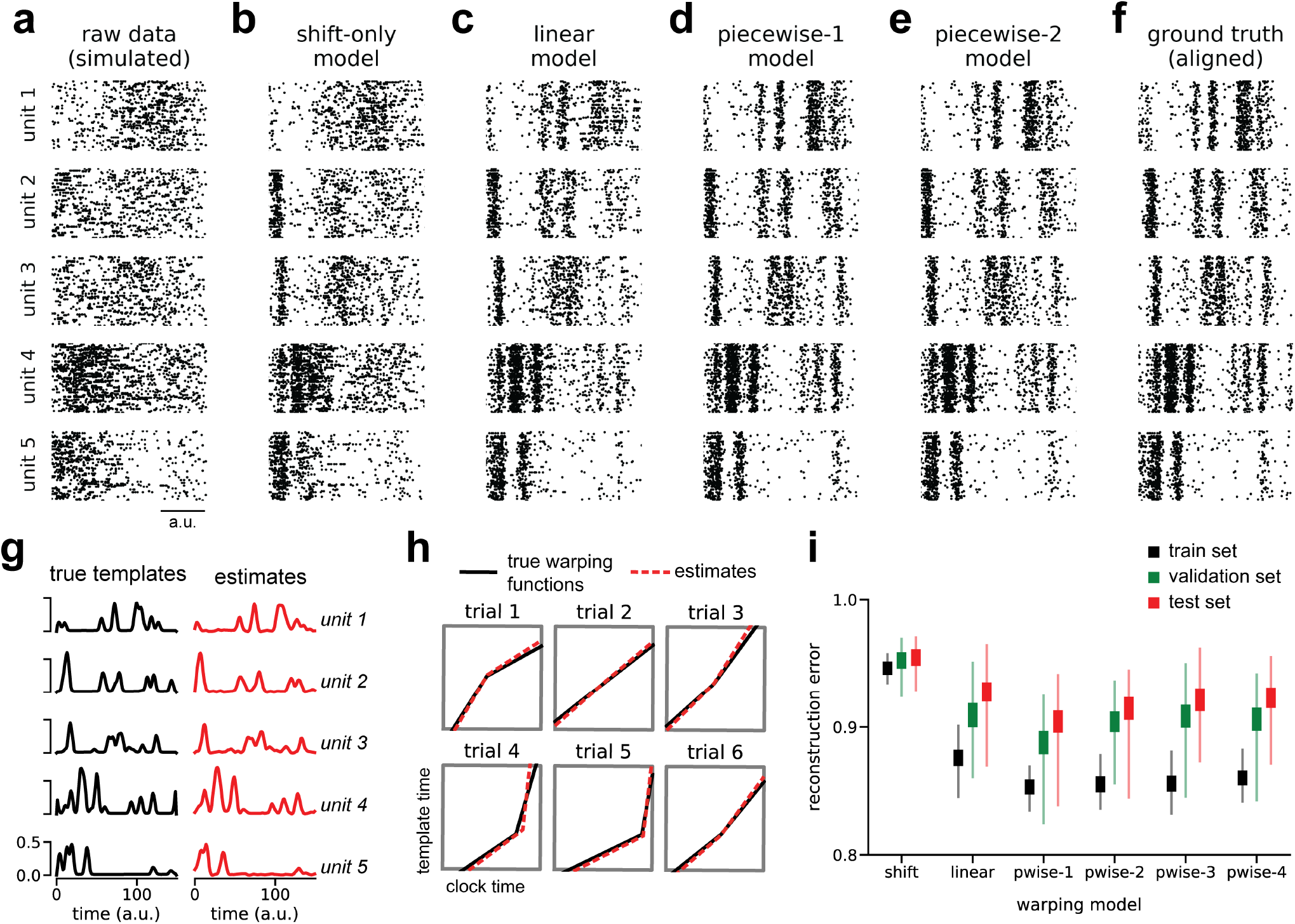
Recovery of ground truth warping functions in synthetic data. A) Synthetic spiking data from *N* = 5 units, *T* = 150 timebins, and *K* = 75 trials. Data were simulated from a ground truth model with piecewise linear warping functions with 1 knot. (B) Data re-aligned by a shift-only warping model. Re-alignment is implemented by applying the inverse warping transformation to the time base of each trial. (C) Data re-aligned by a linear warping model. (D) Data re-aligned by a piecewise-linear (1-knot) warping model. (E) Data re-aligned by a piecewise-linear (2-knots) warping model. (F) Data re-aligned by the ground truth model. Note similarity with panels D & E. (G) Ground truth neural response templates (black) and estimated response templates (red) from the piecewise-linear (1-knot) model. Y-axis denotes the probability of spiking in each time bin. (H) Ground truth warping functions (black) on six representative trials, and estimated warping functions (red) from the piecewise-linear (1-knot) model. (I) Normalized reconstruction errors (norm of residuals divided by norm of raw data) in training, validation, and test partitions of various warping models. Thin lines represent maximum and minimum values; thick lines represent mean *±* standard error. Results were computed over 40 randomized cross-validation runs.

Given these noisy observations, time warping successfully reveals the spike patterns corresponding to the ground truth process. **Figures 2b-e** show model-aligned spike trains (as in **Fig 1f**), across warping models of increasing complexity. These model-derived alignments can be compared to the ground truth spike times after omitting temporal variability from the simulation (**Fig 2f**). The patterns evident in the ground truth data are partially revealed by shift-only and linear time warping (**Fig 2b-c**), but these models are too simplistic to capture the complete fine-scale temporal structure in the data. A piecewise linear warping model with one knot (*piecewise-1* model; **Fig 2d**) accurately captures these details, and represents a parsimonious and “correct” model since it matches the data generation process. Furthermore, the parameters this model closely matched the ground truth response template (**Fig 2g**) and warping functions (**Fig 2h**). Using a slightly more complex model—a piecewise linear model with 2 knots—did not result in substantial overfitting and indeed closely matched the result of the piecewise-1 model (**Fig 2e**).

Identifying a parsimonious warping model is challenging in real-world applications where there is no observable ground truth. To select the appropriate model and regularization strengths we developed a nested cross-validation scheme (see *Methods*). Briefly, we fit the neural response template using a subset of trials, and we fit the warping functions using a subset of the neurons in the data set (the *training set*). For each warping model class (shift-only, linear, piecewise linear, etc.) we select regularization parameters based on a different subset of neurons and trials (the *validation set*). Finally, we evaluate and compare model performance on the remaining data (the *test set*). This procedure is then repeated many times with different partitions of the data. On simulated data, this procedure correctly identifies the piecewise-1 model as having minimal average test error (**Fig 2i**).

In the following sections, we examine the utility of time warping on real neural datasets derived from a variety of sensory and motor areas. The dynamics of these circuits is thought to be closely time-locked to behaviors and sensory cues, yet we found time warping revealed additional temporal structure and precision in all cases, and even identified unexpected oscillatory patterns in two datasets that were decoupled from measured behaviors. Furthermore, we show that simple time warping models (linear or shift-only) are often sufficient to extract these insights, obviating the need for complex, nonlinear warping methods.

### Alignment of olfactory responses to sniff cycle

Mitral/tufted cells in the mouse olfactory bulb display highly variable firing patterns across trials when naïvely aligned to odor delivery (**Fig 3a**). This variability largely stems from trial-to-trial variability in the latency between odor delivery and inhalation, which controls the access of odorants to receptors. Aligning spike times on each trial to inhalation onset reveals a drastically more reliable encoding of the olfactory stimulus (Shusterman et al. 2011).

**Fig 3.**
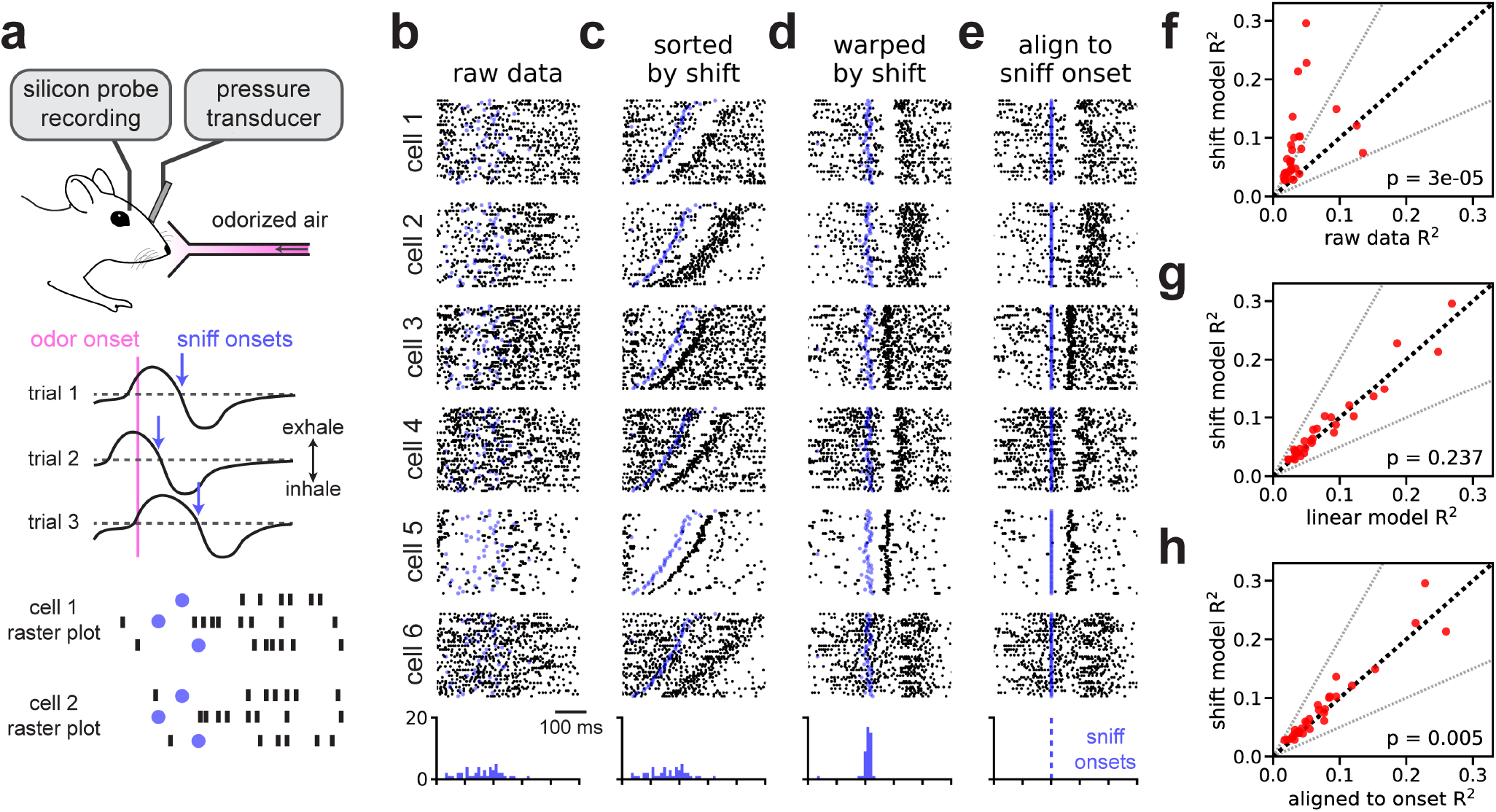
Time warping of mitral/tufted cell activity recovers sniff-locked activity patterns. (A) A head-fixed mouse sampled odorized air (*α*-pinene, 10^−2^ M) while spiking activity of isolated mitral/tufted cells neural activity was recorded. Airflow was switched from a non-odorized source to an odorized source on each trial. Variability in inhalation onset from trial-to-trial caused jitter in the onset of the olfactory response. (B) Spike raster plots for six representative cells over all *K* = 45 trials with spike times aligned to odor delivery. Black dots denote spike times, blue dots denote sniff onset. Blue histogram at the bottom indicates distribution of sniff onset times. (C) Same plots as shown in panel B, but with trials re-ordered by the magnitude of the warping model. (D) Raster plots with spike and sniff onset times re-aligned by applying the inverse warping functions on each trial. (E) Raster plots with spike re-aligned to sniff onset. (F) Trial-to-trial reliability (*R*^2^) for all *N* = 30 cells before (x-axis) and after (y-axis) alignment by the shift-only model. Dashed black line indicates the identity line. Dashed grey lines indicate a two-fold increase or decrease in *R*^2^. The p-value is computed using a Wilcoxon signed rank test. (G) Same as F, but comparing the alignment of the linear warping model (x-axis) to a shift-only warping model (y-axis). (H) Same as F, but comparing the alignment of the shift-only warping model (y-axis) to the data manually aligned to sniff onset (x-axis).

We reasoned that simple time warping models could be used to accurately align mitral/tufted cell activity using purely neural activity, bypassing the need to measure inhalation directly. We tested this hypothesis on a multielectrode recording from *N* = 30 neurons over *K* = 45 trials of odor presentation at a fixed concentration (*α*-pinene, 10^*−*2^M). We found comparable results on a separate set of trials on which a different odorant was presented (**Fig 3, Supplement 1**; limonene, 10^*−*2^ M). We experimentally measured intra-nasal pressure to detect sniff onset and offset. Critically, sniff measurements were not provided to the model and spike times were instead aligned to odor presentation. As expected, this initial alignment strategy produced highly disordered spike rasters for individual neurons (**Fig 3b**).

We found that a shift-only time warping model captured precise sensory responses from these raw data, as revealed by re-sorting the trials based on the model’s shifts (**Fig 3c**) or by applying these shifts to align the raw spike times (**Fig 3d**). Here, as well in all subsequent results, we adopted a leave-one-out validation procedure such that model-aligned spike rasters were computed only for held out neurons. That is, we excluded each cell (1-5 in **Figure 3**) from model fits, and then simply applied each trial’s inverse warping function to the held out cell. Thus any temporal structure seen in **Figure 3d** is unlikely to arise as an artifact of overfitting.

As expected, aligning spike times to inhalation onset times reveals similar patterns in these data (**Fig 3e**). Indeed, the shift parameters learned by the model correlated very tightly with the onset of sniffing (see blue dots and histograms, **Fig 3b-e**). These results are nonetheless a useful demonstration, since the model inferred these precise responses from the neural data alone and without reference to intra-nasal pressure. Furthermore, closer examination suggested that the unsupervised, shift-only model may enjoy slight performance advantages relative to the simple supervised alignment method. For example, when aligned to sniff onset, cells 4 and 5 in **Figure 3** exhibit subtle, but perceptible, jitter in their responses (**Fig 3e**), and this variability is visibly corrected by time warping (compare to **Fig 3d**).

We quantified the trial-to-trial variability of each neuron by computing the coefficient of determination (*R*^2^) of the neuron’s PSTH. In an approach similar to leave-one-out cross-validation, we fit time warping models while holding out neurons one at a time; the *R*^2^ was then computed on the held out neuron before and after warping. Relative to the raw spike times (i.e. odor onset aligned), shift-only time warping improved *R*^2^ in nearly all neurons, with many increasing over two-fold (**Fig 3f**; average 107% increase in *R*^2^, geometric mean; Wilcoxon signed rank test,
*p* < 10^−4^, *n* = 30). Moreover, moving from a shift-only warping model to a more flexible linear warping model did not produce any significant improvements in *R*^2^(**Fig 3g**). Relative to sniff onset alignment, shift-only time warping improved the *R*^2^ criterion mildly (**Fig 3h**; average 11% increase in *R*^2^, geometric mean; Wilcoxon signed rank test, *p* = 0.005, *n* = 30).

### Alignment of motor cortex dynamics during reaching in nonhuman primates

Neural dynamics underlying motor control also exhibit variable time courses due to trial-to-trial differences in reaction times and muscle kinematics. To investigate the benefits of time warping in this setting, we first examined data from a canonical reaching experiment in a nonhuman primate (**Fig 4a**). On each trial, the subject (Monkey J) moved its arm to one of several possible target locations after a mandatory delay period that randomly varied between 300 and 700 ms. In addition to this inherent timing variability due to task design, the monkey exhibited variable reaction time ranging from 293-442 ms (5th and 95 percentiles). We limited our analysis to upward reaches (90° from center) with the target placed at 40, 80, or 120 cm from the center. We observed similar results on other reach angles (**Fig 4, Supplement 1**), as well as when data was pooled across all reach angles (data not shown). Multiunit activity was collected from *N* = 191 electrodes across two Utah multielectrode arrays placed in primary motor (M1) and premotor (PMd) cortices (see *Methods*).

**Fig 4.**
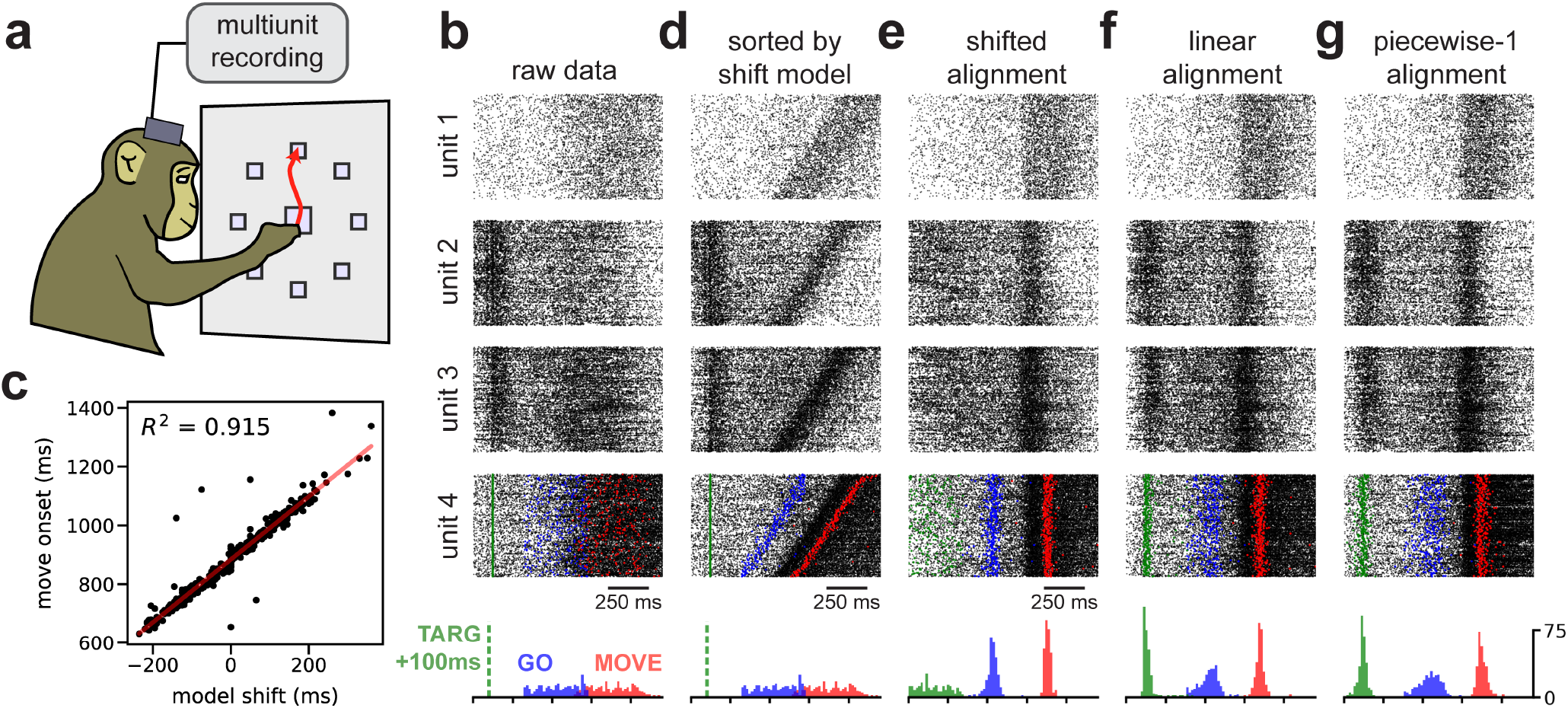
Time warping of reach dynamics in a nonhuman primate. (A) Reaches towards a 90° target were analyzed. (B) Spike data from four example multiunits over all trials. Units 1 & 4 were from primary motor cortex; units 2 & 3 were from pre-motor cortex. At the bottom, the time distribution of task events is shown: 100 ms after target onset (TARG, green), end of delay period (GO, blue), and movement onset (MOVE, red). (C) Scatterplot showing correlation between the hand movement onset on each trial and the learned shift parameter in the shift-only warping model. (D) Same as panel B, but with trials re-sorted by the per-trial shift parameter in the shift-only warping model. (E) Same as panel B, but with spikes aligned according to the shift-only warping model. (F) Same as panel B, but with spikes aligned according to the linear warping model. (G) Same as panel B, but with spikes aligned according to the piecewise-linear warping model with one knot.

The most dramatic changes in neural firing rates are closely time-locked to movement (Churchland et al. 2012; Kaufman et al. 2016). Thus, it is common to track hand position on a moment-by-moment basis and use these measurements to align spike times to the onset of movement or the peak hand velocity on each trial. We instead examined spike trains aligned to the beginning of the delay period (**Fig 4b**), and used time warping to infer an alignment without any reference to the animal’s behavior.

As expected, a shift-only warping model closely aligned spike times with the onset of movement. The model’s learned shift parameter on each trial correlated very tightly with movement onset (**Fig 4c**), achieving a comparable level of performance (*R*^2^ = 0.9) to what was recently reported for a complex, nonlinear warping method (Duncker and Sahani 2018). Furthermore, the shift-only warping model enabled the visualization of movement-related firing rate changes in single-neuron rasters, either by re-sorting the trial order of the raw data (**Fig 4d**) or by re-aligning the spike times (**Fig 4e**).

Thus, learning a single per-trial shift was sufficient to align neural spike times to movement, without any reference to hand tracking data. However, shifting spike times in this manner also *destroyed* other structure in the data. Namely, a subset of multiunits, mostly in PMd, showed increased firing around ~100ms into the delay period—i.e. shortly after the reach target was visually presented to the animal (see units 2 & 3 in **Fig 4**). Due to the variable delay between target onset and movement onset, a shift-only warping model is incapable of simultaneously aligning spikes across these two events.

A linear time warping model more appropriately captures this structure in the data. On each trial, the model utilizes its two free parameters—the slope and intercept of the warping function—to precisely align these two task events (**Fig 4f**). Importantly, as in all of our results, the warping model is fit purely to the neural data without any reference to behavior. Thus, these results provide strong evidence, via an unsupervised time warping method, that reliable neural dynamics occur around the time of movement onset and shortly after target onset. Using nested cross-validation, we determined that more complex, piecewise-linear warping functions did not provide large benefits over the linear warping model; however, the linear warping model provided a reproducible benefit over the shift-only warping model (**Fig 4, Supplement 2**).

### Detection of ~13-40 Hz spike-time oscillations in primate pre-motor cortex

Thus far, we have shown that the temporal alignments learned by simple warping models can closely correlate with behaviors (e.g. movement or sniffing) and sensory cues (e.g. reach target presentation). This agreement demonstrates that time warping models can converge to reasonable and human-interpretable solutions, and, conversely, suggests that established alignment practices in these systems are well-justified from a statistical perspective. However, time warping methods can also uncover more subtle and unexpected features in spike train data.

In primate premotor cortex, the local field potential (LFP) shows prominent oscillations in the beta frequency range (13-40 Hz) during movement preparation, which are correlated with spike timing (Murthy and Fetz 1992; Sanes and Donoghue 1993; Reimer and Hatsopoulos 2010). While recent work has elucidated the statistical relationships between LFP and behavior (Khanna and Carmena 2015; Khanna and Carmena 2017; Chandrasekaran et al. 2019), the impact of beta oscillations on population-level spiking activity is still poorly understood. Recent work used a complex, black box model of neural dynamics to detect oscillatory structure in high-dimensional spike trains (Pandarinath et al. 2018). Here, we show that shift-only or linear time warping models can recover similar oscillations, and compactly summarize single-trial variations in oscillation phase and frequency in the warping function parameters.

We examined premotor cortical data collected from two different monkey subjects (Monkey J and Monkey U) performing point-to-point reaches; one animal performed these reaches with an unrestrained hand, while the other used a manipulandum (see *Methods*). The oscillations are strongest during the pre-movement delay period, and thus we first focused on a time window beginning 400 ms prior to and 100 ms after go cue presentation. We found that having a larger number of trials was beneficial, so we pooled trials from all reach angles for this analysis. We analyzed multiunit data (not spike sorted) for each monkey from *N* = 96 electrodes placed in PMd.

No oscillations were visible in pre-movement spike rasters aligned to go cue (**Fig 5a**; data from Monkey U). However, re-aligning these spike trains based on a shift-only warping model revealed oscillations in virtually all multiunits. These oscillations occurred at ~18 Hz in Monkey U (shown in **Fig 5b**) and at ~40 Hz in Monkey J (**Fig 5, Supplement 1**); these results are within previously reported frequency ranges (Murthy and Fetz 1992; Sanes and Donoghue 1993). In Monkey U, these oscillations were more apparent after linear warping (**Fig 5c**), suggesting that the frequency (in addition to the phase) of the oscillations can be variable on each trial. These oscillations were roughly in-phase across multiunits—as a result, averaging spike counts across all multiunits and trials (**Fig 5a-c**, bottom) produced a cleaner ~18 Hz oscillation in time warped spike trains.

**Fig 5.**
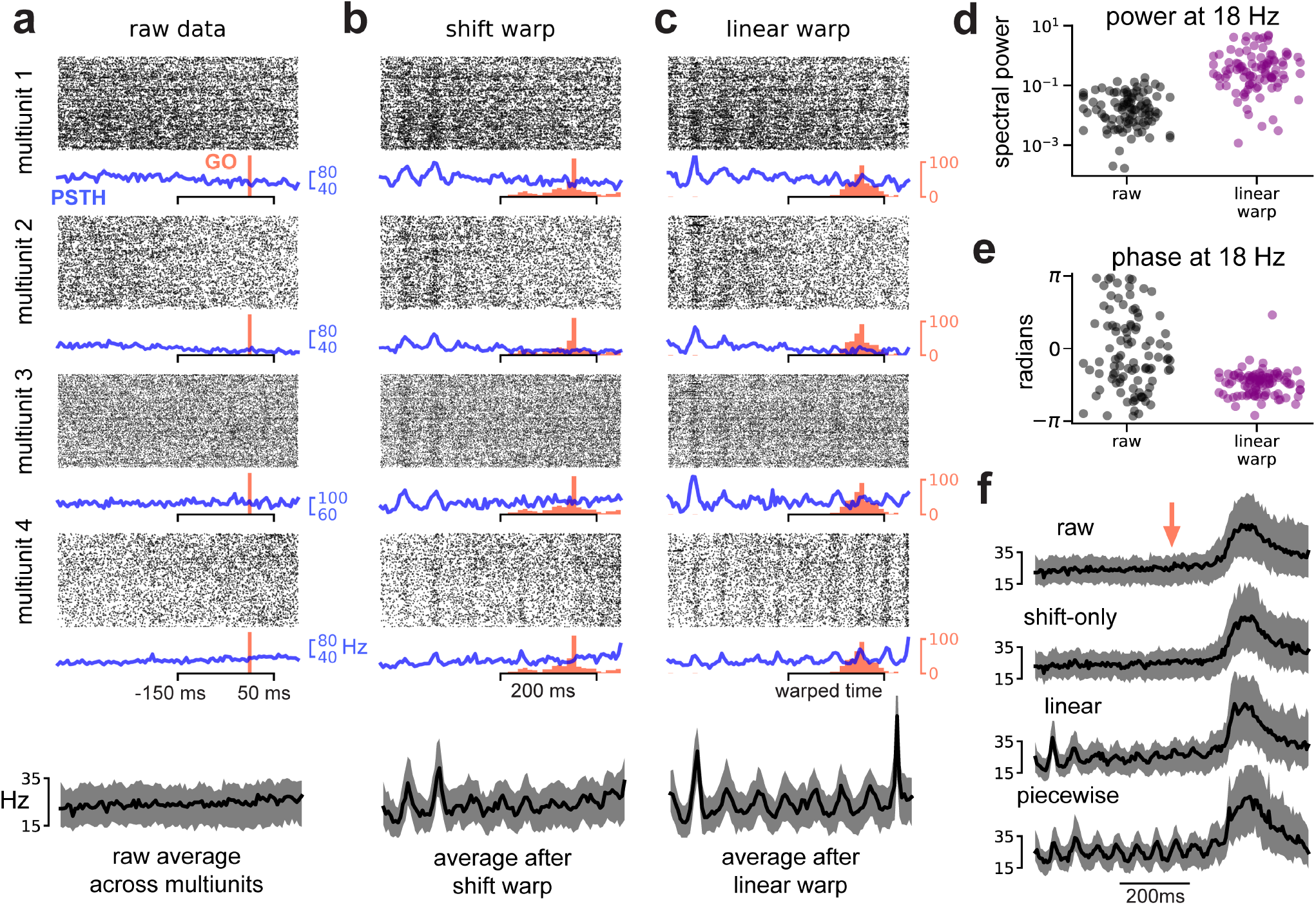
Spike-level oscillations in primate premotor cortex. (A) Multiunit activity aligned to go cue. Four representative multiunits are shown; spike rasters (black) trial-average PSTH (blue) are shown for each multiunit. Go cue onset is shown in red. The average firing rate across all multiunits is shown on the bottom (black trace). Shaded grey region denotes upper and lower quartiles. (B) Same as (A) but with spike times aligned by shift-only warping model. Each displayed multiunit was held out from the model fit. (C) Same as (A) but with spike times aligned by linear warping model. Each displayed multiunit was held out from the model fit. (D) Oscillatory power at 18Hz in trial-average multiunit activity aligned to go cue (black dots) and after linear warping (purple dots). Each dot represents one of *N* = 96 multiunits. (E) Same as (E) but showing oscillation phase for each multiunit. (F) Multiunit activity averaged across electrodes and trials in a larger time window (800 ms) around go cue (red arrow). Oscillations are not visible in the raw data (top) or after shift warping (second from top). Oscillations are recovered by either linear (second from bottom) or piecewise linear warping (bottom). Shaded grey region denotes upper and lower quartiles.

We confirmed that the spike-level oscillations were in-phase with LFP oscillations in Monkey U. To do this, we applied the time warping models fit on spike train data to bandpass-filtered LFP signals (10-30 Hz). The LFP signal was misaligned across trials in raw data, but was accurately aligned by the spike-level time warping models (**Fig 5, Supplement 2**), suggesting that the two signals are coherently time-warped (in this case, temporally shifted and/or stretched) on a trial-by-trial basis. On a methodological level, this demonstrates that time warping models can *generalize* and make accurate predictions about other time series (e.g., LFP) with qualitatively distinct statistics from the training data (e.g., spike times). This ability to identify structure across different data streams in a flexible, unsupervised manner is an attractive feature of time warping models, which is facilitated by our choice to use simple and invertible warping functions.

To quantify these effects more carefully across all multiunits, we compared the PSTHs computed from raw data (blue traces; **Fig 5a**) to PSTHs computed from data aligned by linear time warping (blue traces; **Fig 5c**). The raw PSTHs exhibited no oscillations, as this pattern was temporally jittered and stretched from trial-to-trial and therefore abolished by trial averaging. In contrast, oscillations can be observed to varying extents in the PSTHs computed after alignment by linear warping (which corrects for these trial-to-trial variations). Using Fourier analysis to estimate the amplitude and phase of the oscillation at 18 Hz, we found that alignment by linear warping increased the strength of the oscillation by 1-2 orders of magnitude in most multiunits (**Fig 5d**). Furthermore, in the raw PSTHs the oscillation phases were widely spread across multiunits, consistent with there being no detectable oscillations above background noise (**Fig 5e**; gray dots); in the aligned PSTHs, the phases were tightly clustered, reflecting that nearly all multiunits oscillated in a coherent and detectable manner (**Fig 5e**; purple dots).

We wondered whether time warping would fail to recover these oscillations if the movement-related spiking, which occurs at a much higher firing rate than pre-movement activity, was included in the analysis. To examine this, we fit warping models to a larger time window (±400 ms around go cue presentation), which included the movement-related increase in firing rate. Time warping was still able to extract oscillations under these more challenging circumstances (**Fig 5f**). Interestingly, a shift-only model was no longer sufficient to reliably capture oscillatory activity, suggesting that the oscillations were not phase-aligned with movement onset on a trial-by-trial basis. In contrast, linear or piecewise linear warping functions were able to recover the oscillatory structure (**Fig 5f**; bottom). Thus, while the shift-only model is simplest to interpret, it may be insufficient to capture certain results under particular circumstances. This emphasizes the utility of conceptualizing time warping as a range of models (as in **Figure 1b**) rather than a single method—one can systematically increase the warping complexity to capture increasingly complex features in neural data.

### Detection of ~6-7 Hz oscillations in rat motor cortex

We have seen that time warping can reveal interpretable structure, even under very simple and well-controlled experimental conditions. Discrete reaching, for example, is arguably the simplest volitional motor behavior that one can study, and yet straightforward behavioral alignments obscure salient spike-time oscillations (see **Fig 5**). To study a more complex behavior, in a different animal model, we analyzed motor cortical activity in rats trained to produce a timed motor sequence (Kawai et al. 2015; Dhawale et al. 2017). Rats were trained to press a lever twice with a target time interval of 780 ms, and were rewarded if the sequence was completed within ±80 ms of this target (**Fig 6a**). While rats produce stereotyped motor sequences in this setting, the duration between lever presses and the timing of intermediate motor actions is variable from trial-to-trial. We examined a dataset consisting of *N* = 30 neurons and *K* = 1265 trials; the interval between lever presses ranged from 521-976 ms (5th- and 95th-percentiles) across trials.

This experiment has three obvious alignment procedures: align spike times to the first lever press, align spike times to the second lever press, or linearly stretch/compress the spike times to align both lever presses across trials (i.e. human-supervised time warping). **Figure 6b-d** shows the activity of six example neurons under these alignment strategies. At a high level, these rasters demonstrate that neurons preferentially respond to different behavioral events within a trial. For example, cell 1 in **Figure 6** fires after the second lever press, while cell 6 in **Figure 6** fires after the first press. Thus, it is not obvious which alignment is preferable and indeed different insights may be gained from analyzing each.

Unsupervised time warping revealed structure in the data that is hidden in all three behavioral alignments. A shift-only warping model uncovered strong oscillations in many neurons, as visualized either by re-sorting trials based on the learned shift (**Fig 6e**, same alignment as **Fig 6b**), or by using the model to re-align spike times (**Fig 6f**). These findings are not due to spurious alignments produced by an overfit model. Each spike raster in **Figure 6f** was generated from data held-out from the model—that is, the visualized cells were excluded during optimization of the warping functions, and the learned alignment transformation was then applied to this held out data.

**Fig 6.**
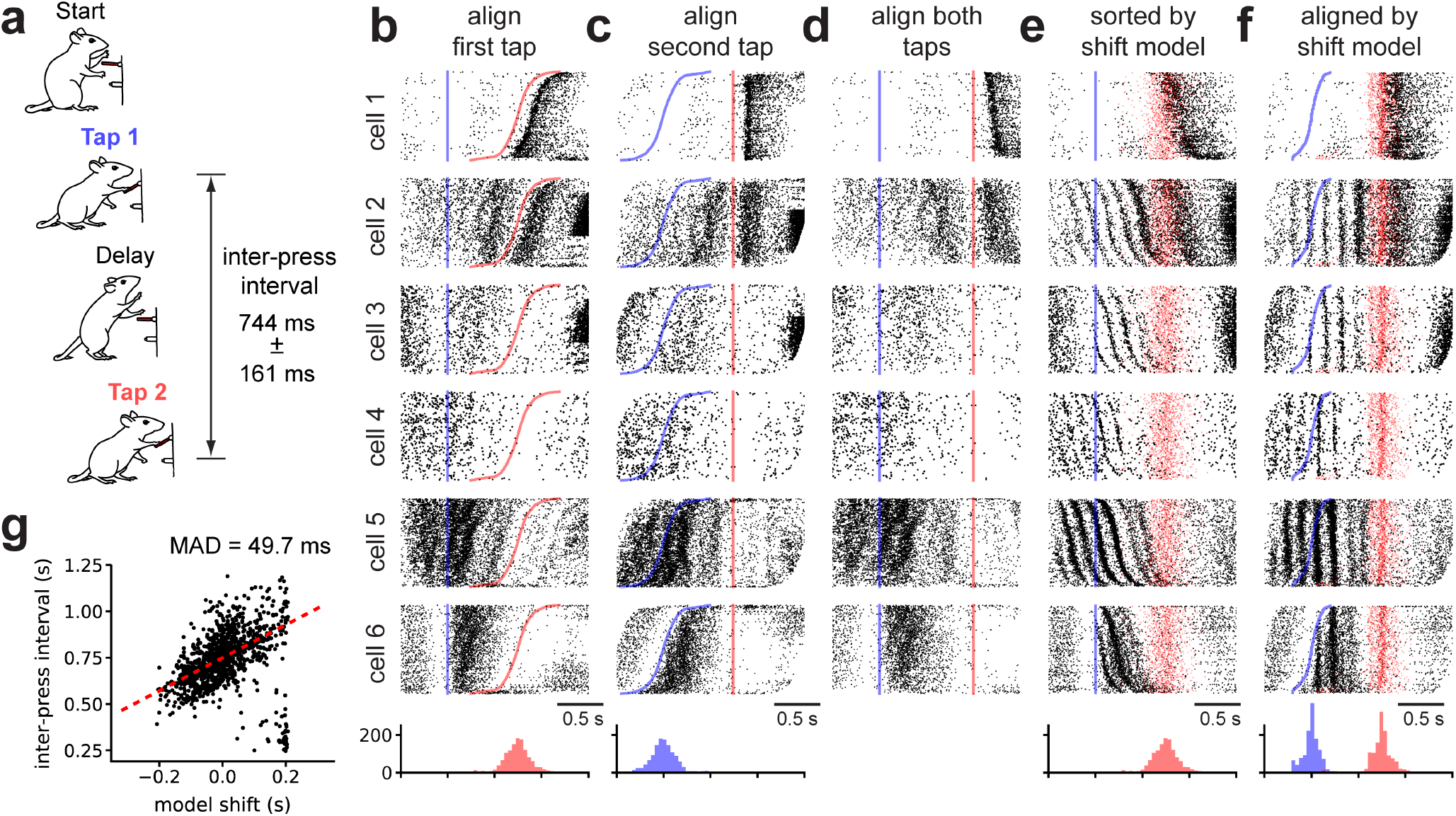
Shift-only time warping reveals temporally precise theta-locked oscillations in rat motor cortex. (A) Rats were trained to press a lever twice with a prescribed temporal delay. The median *±* IQR inter-press-interval is listed. (B) Spike raster plots for six representative cells over all trials with spike times aligned to the first lever press (blue line) and trials sorted by the inter-press-interval. The time of the second lever press is denoted by the red line in all plots. Red histogram at the bottom denotes the distribution of the second lever press times. (C) Spike raster plots re-aligned to the second lever press. Blue histogram at the bottom denotes the distribution of the first lever press times. (D) Spike raster plots aligned by linearly stretching/compressing the time axis in each trial so that the first and second lever presses were both aligned (note the lack of scale bar, as time is no longer constant across trials). (E) Same spike rasters as in panel B, but with trials re-sorted by the magnitude of a shift-only warping model. The time of the second lever press on each trial is denoted by a semi-transparent red dot. (F) Spike raster plots after re-sorting trials as in panel E and shifting the spike trains on each by the shift-only time warping model. (G) Relationship between per-trial shift learned by the shift-only time warping model (horizontal axis) and measured inter-press interval (vertical axis) on each trial. No tight correlation with behavior is observed—dashed red line denotes a robust linear regression fit (Huber loss function, ϵ = 1.001), and the Median Absolute Deviation (MAD) is listed as a measure of fit.

These results reveal a partial decoupling of behavioral events (lever presses) with neural firing patterns. After alignment, both the first and second lever presses occur at variable times within each trial (**Fig 6f**, histograms at bottom). Furthermore, the learned shift on each trial only loosely correlated with inter-press interval (**Fig 6g**). Taken together, these features of the data suggest that it would be difficult to discover this oscillatory structure by manual alignment, demonstrating the power of unsupervised time warping models.

While the uncovered oscillations are not phase locked with lever press times, they are nonetheless correlated with certain aspects of the animal’s behavior. In particular, some cells only exhibit the ~6-7 Hz oscillation following the first lever press, with remarkable temporal precision (see cells 3 and 6 in **Fig 6f**). Indeed, multiple cells exhibit non-oscillatory firing prior to the first lever press, but rapidly switch to an oscillatory behavior following the lever press (see cell 3 in **Fig 6f** and **Fig 6, Supplement 1**). Other cells exhibit oscillations prior to the first lever press, but the amplitude and precision of the oscillations appears to improve following the first lever press (see cell 4 in **Fig 5F**). Still other cells either do not exhibit oscillations (cell 1 in **Fig 6f**) or exhibit strong oscillations both prior to and following the first lever press (cell 5 in **Fig 6f**). Time warping enables us to discover and visualize this full spectrum of functional cell types, which are otherwise difficult to detect and characterize. The presence of oscillations in single neurons can be confirmed by plotting the distribution of inter-spike-intervals (**Fig 6, Supplement 1**); the shift-only model goes beyond this method by demonstrating that a large population of neurons are coherently phase-shifted on a trial-by-trial basis, and by enabling characterization of the full population dynamics and behavioral events in an aligned temporal space.

We then examined whether these spike-level oscillations were aligned with oscillations in LFP. The average frequency spectrum of the LFP did display a prominent peak at ~6-7 Hz—a very similar frequency range to the spike-level oscillations identified in **Figure 6**. To characterize the relationship between these two oscillatory signals, we bandpassed filtered the LFP between 5-9 Hz on each trial and fit a separate shift-only time warping model to the LFP traces. The time warping functions learned on LFP data did not uncover the spike-time oscillations shown in **Figure 6**, and likewise the LFP signals were not well-aligned by the time warping functions fit to spike times (**Fig 6, Supplement 1**). Thus, unlike the oscillations identified in primate premotor cortex, the oscillations in rat motor cortex were not aligned with the LFP. This analysis illustrates two useful features of our time warping framework. First, the models can be flexibly applied to other data types beyond spiking data (e.g. LFP). Second, if two or more data types are simultaneously collected (e.g., LFP and spikes), separate time warping models can be fit to each signal and then compared to assess whether these signals are coherently warped on a trial-by-trial basis. These post-hoc comparisons are drastically simplified when time warping functions are constrained to be linear or piecewise linear, instead of fully nonlinear.

## Discussion

While the temporal precision of neural coding has been a matter of intense debate, few studies have leveraged statistical alignment methods to address this problem. Earlier work incorporated time warping into single neuron encoding and decoding models (Aldworth et al. 2005; Gollisch 2006; Smith and Paninski 2013; Lawlor et al. 2018), as well as dimensionality reduction methods (Poole et al. 2017; Duncker and Sahani 2018). Here, we decoupled time warping from these other modeling objectives, to achieve a flexible and simplified framework. We surveyed a broader range of datasets than past work, spanning multiple model organisms, brain areas, and sensory/motor tasks. In all cases, we found that the simplest and most interpretable models—often those with shift-only or linear warping functions—matched the performance of more complex models, while uncovering striking and sometimes unanticipated dynamics.

We first examined two datasets in which behavioral alignments are well-established, and found that unsupervised time warping inferred similar alignments based on neural data alone. In mouse olfaction, mitral/tufted cells exhibit reliable sensory coding when spike times are aligned to sniff onset, but not odor onset (Shusterman et al. 2011). The simplest time warping model—a shift-only model—accurately realigned spikes without any reference to intranasal pressure measurements, and more complex warping methods (linear and piecewise linear) produced little additional benefit. In primates executing cued reaches, a shift-only time warping model accurately predicted movement onsets (*R*^2^ ≈ 0.9) without any reference to hand tracking data. This performance is comparable to recent work that developed a nonlinear warping method (Duncker and Sahani 2018), and surpasses prior work (Petreska et al. 2011; Poole et al. 2017).

Together, these results demonstrate that shift-only and linear warping models can match or even outperform more complex methods. These simpler models have two attractive properties. First, they manipulate model estimates of single-trial firing rates in a more interpretable manner (see **Fig 1**), enabling exploratory data analysis and visualization. Second, we developed fast and computationally scalable optimization methods for this class of models. On a modern laptop, these models can typically be fit to data from 1000 neurons, 100 timepoints, and 1000 trials in one minute or less. This scalability is of great practical importance given the exponentially increasing size of neural recordings (Stevenson and Kording 2011), and the growing need for rigorous cross-validation and model comparison methods (Chandrasekaran et al. 2018), which are often computationally intensive if not prohibitive.

Time warping also uncovered firing patterns that were not aligned to any stimulus or measured behavior. For example, we observed ~13-40Hz spike time oscillations in primate premotor cortex during movement preparation (see **Fig 5**), which we then verified were phase-aligned with LFP (see **Fig 5, Supplement 2**). Notably, the time warping models we used did not assume any oscillatory structure in the data, and thus provide a data-driven validation that spike-level oscillations are a salient feature of the dynamics. Furthermore, a linear time warping considers the activity of the full neural population to estimate the changes in the phase (y-intercept) and frequency (slope) of the oscillation on a trial-by-trial basis. This population-level approach can be contrasted with popular frequency-domain statistical measures like *coherence*, which measures the degree of phase synchronization between two spike trains or between a single spike train and LFP (Fries et al. 1997; Jarvis and Mitra 2001; Sun et al. 2005; Aoi et al. 2015). Future work could incorporate oscillatory basis functions into time warping models to combine the benefits of pairwise spectral analysis with the population-level modeling perspective adopted in this paper. By drawing statistical power from larger numbers of simultaneously recorded neurons, research in this direction could provide tighter links between spike-based and LFP-based measures of oscillation, which have been difficult to characterize despite extensive prior work (Ray 2015).

However, oscillatory patterns may not always be synchronized to LFP or pre-conceived behavioral variables, as we observed in rat motor cortex (see **Fig 6**). While further work is needed to fully elucidate the properties and functions of these ~7 Hz oscillations, we found that they were, in some neurons, gated by a motor action—specifically, the first lever press—suggesting a potential relevance of these oscillations to the motor time keeping task (**Fig 6, Supplement 2**). Another, possibility is that orofacial behaviors such as whisking and licking are the primary driver of these oscillations (Hill et al. 2011). Other work has shown that persistent ~7 Hz LFP oscillations may be locked to the respiration cycle (Tort et al. 2018); the transient, spike-level oscillations we observed were decoupled from LFP, and thus likely distinct from this phenomenon. Regardless of their root cause, this result demonstrates the ability of time warping to extract unexpected features of scientific interest from high-dimensional spike trains. Thus, while it will be interesting to develop specialized extensions to time warping that address particular scientific questions (e.g. oscillatory firing patterns), the general-purpose framework developed here can be a powerful tool for exploratory analysis, as it makes few pre-conceived assumptions about the data.

It is possible that future work using more complex, nonlinear warping methods can uncover even finer structure in neural data. However, we observed that DTW and other classical methods were prone to overfit data, suggesting that careful regularization will be needed for this approach to succeed. A recently proposed method, called *soft-DTW*, looks promising (Cuturi and Blondel 2017). While the method is mathematically elegant, we found that soft-DTW can be difficult to interpret as it does not represent temporal alignments as a single warping function, but rather uses a weighted combination of all possible warping paths. In general, nonlinear warping methods will require careful application, cross-validation, and secondary analyses to be useful statistical tools for neuroscience.

Time warping is only one form of variability exhibited by single-trial neural dynamics. We purposefully examined time warping models in the absence of other modeling assumptions, such as trial-to-trial variation in amplitude (Bollimunta et al. 2007; Goris et al. 2014; Williams et al. 2018), or condition-specific changes in dynamics (Duncker and Sahani 2018). We also made the restrictive assumption that all neurons share the same time warping function on an individual trial (Shokoohi-Yekta et al. 2015). Finally, we assumed and exploited a trial structure to neural time series data throughout this work. To study more unstructured time series, future work could incorporate time warping into state space models (Macke et al. 2015) or sequence extraction algorithms (Mackevicius et al. 2019). Despite these exciting prospects for future statistical methodology, our work demonstrates that even a simple time warping framework can provide a rich and practical set of tools for the modern neuroscientist.

While our results already show that averaging over short, stereotyped trials can obscure fine temporal oscillations and firing events, these shortcomings are undoubtedly more severe in behaviors that have longer temporal extents and exhibit more variability. Thus, we expect time warping methods to play an increasingly crucial role in neural data analysis as the field moves to study more complex and unstructured animal behaviors (e.g. under more naturalistic settings; Krakauer et al. 2017). Furthermore, in complex experimental tasks involving large numbers of conditions and exploratory behaviors, the same motor act or sensory percept may present itself only a small number of times. In this trial-limited regime, precise data alignment may be critical to achieve the necessary statistical power to make scientific claims. We expect simple models, such as linear and piecewise linear warping, to perform best on these emerging datasets due to their interpretability, computational efficiency, and robustness to overfitting.

## Acknowledgements

We thank Isabel Low and Chandramouli Chandrasekaran for feedback and detailed discussions on the manuscript. This work received support from the following research grants and foundations: Department of Energy CSGF fellowshp (A.H.W.), NIH NRSA 1F31NS089376-01 (E.M.T.), Stanford Graduate Fellowship (E.M.T.), NSF IGERT 0734683 (E.M.T.), NIH NINDS T-R01NS076460 (K.V.S.), NIH NIMH T-R01MH09964703 (K.V.S.), NIH Director’s Pioneer Award 8DP1HD075623 (K.V.S.), DARPA BTO “REPAIR” N66001-10-C-2010 and “NeuroFAST” W911NF-14-2-0013 (K.V.S.), the Simons Foundation Collaboration on the Global Brain awards 325380 and 543045 (K.V.S.), ONR N000141812158 (K.V.S.), the Howard Hughes Medical Institute (K.V.S.), Charles A. King Trust (A.K.D.), the Life Sciences Research Foundation (A.K.D.), NIH NINDS R01-NS099323-01 (B.P.O.), NIH R01-DC013797 (D.R.), NIH R01-DC014366 (D.R.), Burroughs Wellcome (S.G.), Alfred P. Sloan Foundation (S.G.), Simons Foundation (S.G.), McKnight Foundation (S.G.), James S. McDonell Foundation (S.G.), and the Office of Naval Research (S.G.).

## Methods

### Code Implementation & Availability

Our code for fitting linear and piecewise linear time warping models is distributed as a GitHub repository (under an MIT license): https://github.com/ahwillia/affinewarp. Our Python implementation relies on the standard SciPy scientific computing libraries (Jones et al. 2001–; Hunter 2007). Additionally, we achieved substantial performance enhancements by leveraging numba, a Python library that enables just-in-time (JIT) compilation (Lam et al. 2015). Step-by-step tutorials for executing our code are available on GitHub.

### Detailed Description of Time Warping

Notation We follow the same notation introduced in the main text. Matrices are denoted in bold, uppercase fonts, e.g. **M**, while vectors are denoted in bold, lowercase fonts, e.g. **v**. Unless otherwise specified, non-boldface letters specify scalar quantities, e.g. *S* or *s*. We use **M**^T^ and **M**^−1^ to denote the transpose and inverse of a matrix, respectively.

We consider a dataset consisting of *N* features over *K* trials with *T* timesteps per trial. For simplicity, we refer to *N* as the number of neurons in the dataset; however, *N* could also refer to the number of fMRI voxels, multiunits, or regions of interest in imaging data. The full dataset is a third-order tensor (a three-dimensional data array) with dimensions *K* × *T* × *N*. The *k*^th^ slice of the data tensor is a *T* × *N* matrix **X**_*k*_, which represents the activity of the neural population on trial *k*. We denote a single element of the tensor as *X*_*k,t,n*_, which specifies the activity of neuron *n* at timebin *t* on trial *k*.

The time warping model produces an estimate of population activity on each trial. Mirroring standard notation in linear regression, we denote the model estimate on trial *k* as 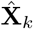 (a *T* × *N* matrix).

### Model Estimate and Template Interpolation Scheme

The main idea behind time warping is to approximate each trial, **X**_*k*_, as a warped version of a *N* × *T* template, 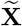, that is shared across all trials. For neuron *n*, at time bin *t*, on trial *k*, the spirit behind the model is:

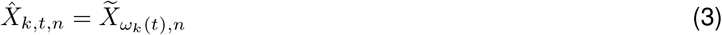

However, this expression is only valid when the warping function, *ω*_*k*_(*t*), produces integer values. To allow the warping functions to produce non-integer values, we adopt a standard linear interpolation scheme. Let *ω*_k_ : *t → τ* describe the time warping function for trial *k*, such that *t* is the integer-valued time index for the data (clock time), and *τ* is any real number representing time for the response template. Then, the model estimate for neuron *n*, at time bin *t*, on trial *k* is given by:

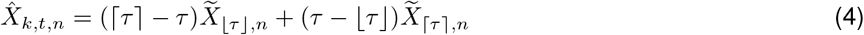

where *τ* = *ω*_*k*_(*t*), ⌈ represents the “flooring” operation, and ⌉ represents the “ceiling” operation. Note that *τ* implicitly depends on the trial index *k*, but we do not explicitly denote this dependence for notational simplicity.

Because the model estimate (**Eq 4**) is a linear combination of 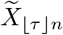 and 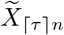, the warping transformation can be represented as a matrix **W** with elements:

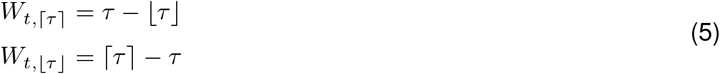

For each trial, the *warping matrix* **W**_*k*_ can be uniquely determined from the warping function *ω*_*k*_. Thus, the model estimate on each trial is given by:

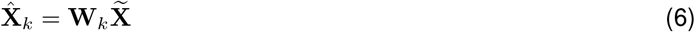

### Optimization Strategy

The model template and warping functions are optimized to minimize an objective function, which we denote as *F* 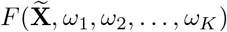. We assume that this objective function decomposes across trials as follows:

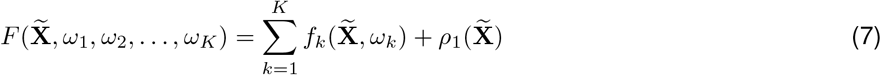

Here *f*_*k*_ is a function defining the model loss on trial *k*, and *ρ*_1_ is a regularization term, penalizing the roughness and size of the template (described in the next section). Our online code package supports least-squares and Poisson loss functions; we adopted the least-squares criterion for the purposes of this paper due to its computational efficiency and its ability to be adapted to non-spike time data (e.g. fMRI or calcium imaging). Under this choice, the per-trial loss function is:

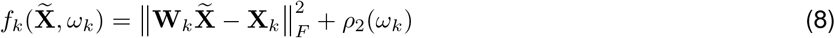

Here, *ρ*_2_ is a regularization term that penalizes the magnitude of warping (described in the next section), and 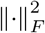 denotes the squared Frobenius norm, which is simply the sum of squared residuals, 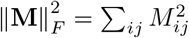.

To minimize *F*, we adopt an alternating optimization (block coordinate descent) approach (Wright 2015). First, each warping function is initialized to be the identity, *ω*_*k*_ (*t*) = *t*, and the template and warping functions are cyclically updated according to the following sequence of optimization subproblems:

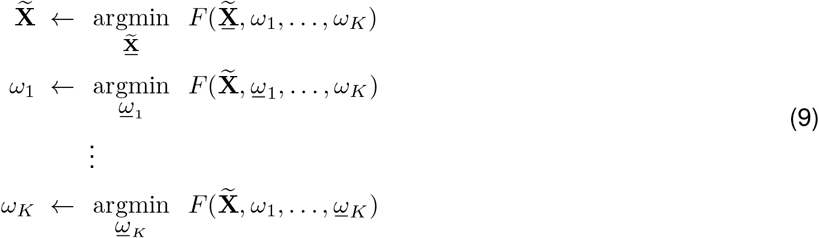

Here, an underlined variable denotes a dummy variable that is optimized over in each subproblem. This sequence of parameter updates is cyclically repeated until the objective value ceases to improve; by construction, the objective monotonically decreases at each step of the algorithm so convergence is guaranteed under mild assumptions (Wright 2015).

This partitioning the parameter updates enables each subproblem to be solved very efficiently. When the template is considered a fixed variable, the objective function decouples across trials (**Eq 7**), which simplifies the warping function updates considerably:

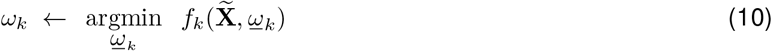

These parameter updates are entirely independent, with each update only depending on the raw data for trial *k*, **X**_*k*_, and the current warping template 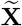. Our code package executes them efficiently in parallel across CPU threads. Furthermore, each warping function is controlled by a small number of parameters in our framework—at best a single parameter (shift-only warping) and at worst only a few parameters (piecewise linear warping). Thus, we perform these updates by a brute force random search (see *Warping Function Regularization and Update Rule*).

The response template is also very simple to update, especially under a least-squares loss criterion. Assume for the moment that the model is not regularized; i.e., 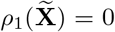 and *ρ*_2_(**W**_*k*_) = 0. Then, because each **W***k* is held constant, updating the template amounts to a least-squares problem that can be solved in closed form:

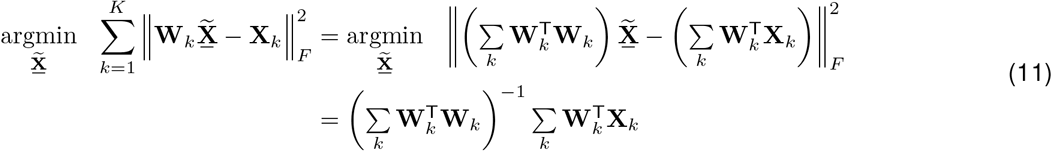

Furthermore the matrix 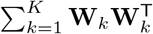 is a *symmetric, tridiagonal matrix*. Intuitively, the tridiagonal structure arises from the constraint that each warping function is monotonically increasing, and the local structure of the linear interpolation scheme. Consider any warping matrix **W**, with an associated warping function *ω*. **Equation 4** implies that *W*_*i,t*_*W*_*j,t*_ = 0 if |*ω*(*t*) − i| > 1 or if |*ω*(*t*) − *j*| > 1 and thus [**W**^*T*^**W**]_*i,j*_ = 0 if |*i* − *j*| > 1.

The tridiagonal structure of 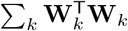 enables the template parameters to be updated extraordinarily fast for practical applications. We use a specialized solver for systems of linear equations with banded, symmetric structure (scipy.linalg.solveh banded). This allows the matrix inversion in **Equation 11** to be (implicitly) carried out in *O*(*T N*) operations instead of *O*(*T*^3^ + *T*^2^ *N*) operations if **W** was treated as a dense matrix.

### Template Regularization and Update Rule

We found that introducing regularization (penalties on the magnitude or complexity of model parameters) can improve the interpretability of the model and its ability to predict held out data. First, we found in some datasets that the warping template could exhibit rapid, high-frequency changes in firing rate (see, e.g., the template in **Fig 1C**, which was fit without regularization). These irregularities likely correspond to the model overfitting to noisy neuronal data, and can be discouraged by penalizing the magnitude of the second finite differences along the temporal dimension of the template (Grosenick et al. 2013; Maheswaranathan et al. 2018). We refer to this term as a *roughness penalty* or *smoothness regularization*. Second, it is possible that the matrix 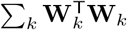 appearing in eq. (11) would become non-invertible or ill-conditioned during optimization. To prevent this, and to discourage the template firing rates from becoming too large, we added a penalty on the squared Frobenius norm of the template. Formally, the regularization on the template is given by:

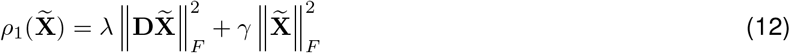

where *λ* > 0 controls the strength of the roughness penalty and *γ* > 0 controls the strength of the Frobenius norm penalty. The matrix **D** is a (*T* − 2) × *T* matrix that computes second-order finite differences:

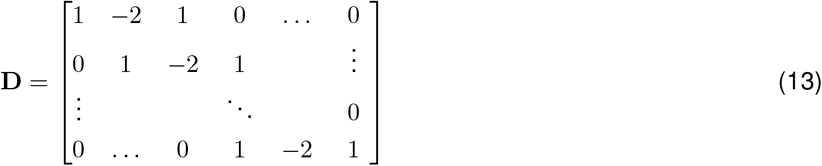

Incorporating this regularization term into the update of the warping template (eq. (11)), we get:

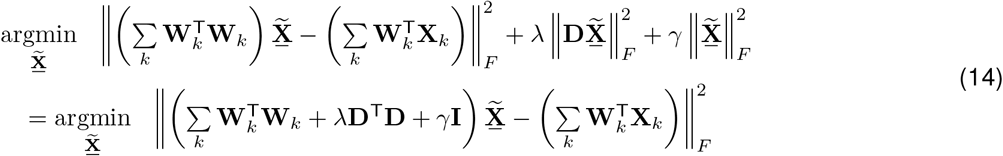

Which yields the template update rule:

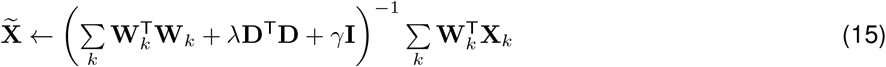

Thus, the solution is the same as before except a term *λ***D**^T^**D** + *γ***I** is added to the inverted matrix (left-hand side of linear system). These modifications hardly affect the computational complexity of the parameter update since *λ***D**^T^**D** + *γ***I** is also a symmetric, banded matrix. Furthermore, as long as *γ* > 0 the overall matrix is positive definite and therefore guaranteed to be invertible.

In practice, we have found that it is simple to hand-tune the regularization strengths for exploratory analysis (though cross-validation procedures, described below, should always be used to monitor for overfitting). We typically set the L2 regularization (*γ*) to be zero or very small (e.g., 1e-4) and do not tune it further. A reasonable value for the roughness penalty scale can be found by visually inspecting the template for various neurons (columns of 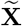) and increasing λ if these time series appear noisy.

### Warping Function Regularization and Update Rule

We found that the optimization landscape of linear and piecewise linear warping functions is complex and full of local minima. Thus, gradient-based optimization methods can be ineffective. Thankfully, the warping functions are (a) low-dimensional and (b) entirely decoupled across trials. Thus, when updating the warping functions, we perform a brute force parameter search for each trial in parallel. For shift-only warping models, we perform a dense grid search over the parameter (the magnitude of the shift).

For piecewise linear warping models we perform an annealed random search as follows. Consider a warping function *ω*(*t*) for any arbitrary trial (we drop the trial index *k* for brevity). We parameterize the warping function as:

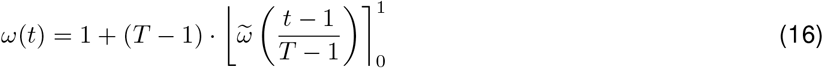

where 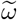 is a piecewise linear function mapping the unit interval [0, 1] to any real number, and 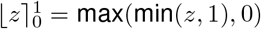 denotes clipping any real number *z* to have a value between zero and one.

The piecewise linear function 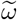 is defined by a series of *M* x-y coordinates, {(*α*_1_, *β*_1_), (*α*_2_, *β*_2_), … (*α*_*M*_, *β*_*M*_)}, where 0 = *α*_1_ < *α*_2_ < 2026;< *α*_*M*_ = 1 and *β*_1_ ≤ *β*_2_ ≤ … ≤ *β*_*M*_. We refer to these coordinates as the *knots* of the warping function. The function is defined using linear interpolation:

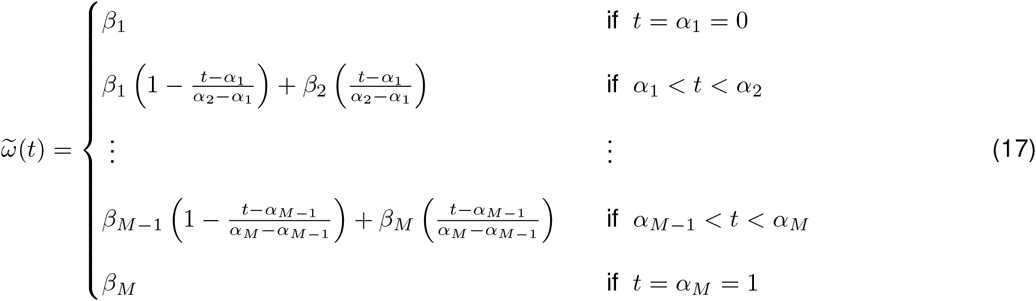

To optimize the warping functions we perform a random search over these coordinates/knots. Let ***α*** = [*α*_1_, *α*_2_, …, *α*_*M*_] and ***β*** = [*β*_1_, *β*_2_, …, *β*_*M*_] denote the current coordinates. We form a new proposed warping function by:

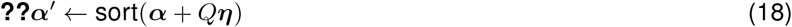

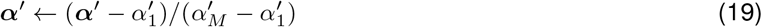

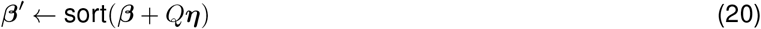

where *Q* > 0 is a scalar parameter tuning the amount of exploration, and ***η*** is a vector of random normal variables with mean zero and unit variance. The procedure “sort(**v**)” re-orders the elements of a vector so that they are in ascending order. If the proposed warping function improves the objective function, we accept the new parameters:

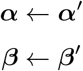

For each round of optimization we exponentially relax *Q* from 1.0 to 0.01 over a fixed number of iterations. We also found that penalizing the warping functions based on their distance from the identity line was helpful in some cases. Intuitively, this encourages the warping functions to be minimal—as the penalty strength increases the warping functions will approach *ω*(*t*) = *t*, resulting in no warping at all in this extreme limit. Similar penalties or hard constraints on time warping have been examined in prior literature (see e.g., Zhang et al. 2017). We chose the penalty to be the area between the unclipped warping function and the identity line:

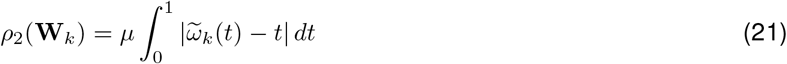

which, for piecewise linear functions with relatively small *M*, can be efficiently computed as the sum of triangular and trapezoidal regions. Here, *μ* ≥ 0 is a scalar hyperparameter controlling the strength of the penalty. In practice we start with *μ* = 0 and increase it if, upon visual inspection, the warping functions are highly deviant from the identity line. Increasing *μ* in these cases can result in more sensible and interpretable templates. Again, cross-validation procedures can be used to asses whether *μ* is too low (resulting in overfitting) or too high (resulting in underfitting).

### Cross-validation

As with any statistical method, one must be very careful that time warping does not reveal spurious structure and features of the data. In **Fig 1**, we saw that even a simple linear warping model can result in noticeable overfitting on a simple synthetic time series. An important technical contribution of our work is a rigorous cross-validation framework for time warping models. This framework, described in detail below, enables us to fine-tune all regularization terms—i.e. the hyperparameters {*γ*, *λ*, *μ*}—across all warping models. That is, we can rigorously compare the performance of shift-only, linear, and piecewise-linear time warping models on an even footing, and thus critically examine the degree of nonlinearity in time warping. While cross-validation is a common procedure in statistical modeling and in modern neuroscience, there are subtle pitfalls that must be avoided in unsupervised learning models (Bro et al. 2008; Perry 2009), and in models with smoothness regularization terms (Opsomer et al. 2001). These concerns are not merely theoretical; they have directly impacted recent results in neuroscience (Latimer 2018).

To properly compare the performance of different warping models, it is important to perform *nested* cross-validation, so that regularization terms are separately tuned for each model. For example, a piecewise linear warping model will often require stronger smoothness and warp regularization terms, compared to a simpler, shift-only warping model. Thus, on each cross-validation run we split the data in three partitions: a training set, a validation set, and a test set. For each model class (shift-only, linear warping, piecewise-linear warping, etc.) we fit 100 models with randomized regularization strengths to the training set; we then evaluated all 100 models on the held-out validation set; finally, the best-performing model was evaluated on the test set. The test set performance is then compared across model classes. We used ~73% of the data for training, ~13% for validation, and the final ~13% for testing. We performed this overall cross-validation procedure 100 times, drawing different randomized data partitions each time—this is known as *randomized cross-validation* and is useful for heterogeneous datasets, in which features (i.e. neurons) exhibit varied levels of noise.

Recall that our dataset consists of *N* neurons, *T* timebins, and *K* trials. The question then arises, should we hold out neurons, timepoints, or trials during cross-validation? Since the warping functions are assumed to be shared across all neurons, these model parameters can be fit on a subset of neurons (training set), and then evaluated on held out neurons (validation/test sets). However, if we hold out individual neurons entirely, then it is impossible to fit the response template matrix 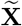 for those cells. Conversely, the response template can be fit to a subset of trials (training set) and evaluated on the remaining trials (validation/test sets). However, if we hold out individual trials entirely, then it is impossible to fit the warping functions associated with those held out trials.

To circumvent this problem we adopt a bi-cross-validation hold out pattern (Owen and Perry 2009). This entails separately and independently partitioning neurons and trials. Thus, we randomly choose training neurons (~73% of all cells), validation neurons (~13% of all cells), and testing neurons (the remaining ~13%). Additionally, we randomly choose training trials, validation trials, and testing trials, according to these same ratios. The model warping functions are fit to all trials, but only on the training neurons; the response template is fit for all neurons, but only on the training trials. When reporting the training performance, we compute the reconstruction loss on the *intersection* of the training neurons and training trials. Likewise, when evaluating models on the validation (or test) set, we compute the the reconstruction loss on the intersection of the validation (or test) neurons and validiation (or test) trials.

Temporal dependencies in model errors can complicate proper cross-validation (Opsomer et al. 2001; Latimer 2018). To avoid these complications, we leave out entire trials for individual neurons, rather than leaving out a subset of time bins.

### Null models and other sanity checks

The cross-validation procedure described above is fully rigorous, but computationally expensive to perform. Even if each optimization run only takes a few seconds to complete, comparing *M* warping models over *P* random samples of the regularization parameters, and repeating the whole process over *Q* randomized folds leads to long run times; for example, ~26 hours for *M* = 5, *P* = 100, *Q* = 50 and each model taking ~20 seconds to optimize. This rather unfavorable scaling underscores why our attention to performance enhancements—e.g. by exploiting banded matrix structure when updating the model template—is critical for practical applications. On the other hand, a full cross-validation run is often unnecessary for exploratory data analysis and visualization. Here we briefly outline two simple procedures for validating time warping visualizations in a more interactive manner. Our online code package also supports both of these options.

First, one can create a very simple null dataset of neural activity that, by construction, contains no warping. By comparing the results of time warping on this null dataset to those achieved on the real data, we gain an informative reference point. For spiking data, we simulate null data by computing the trial-average firing rate of each neuron and then drawing Poisson i.i.d. random spike trains on every trial. That is, on each trial, the spike train for a neuron is drawn from an inhomogeneous Poisson process, with a rate function given by the trial-average firing rate. Similar baselines could be developed for calcium imaging and fMRI studies after specifying an appropriate noise model.

Second, a key visualization tool enabled by time warping is the alignment of neural activity across trials. This alignment is achieved by applying the inverse warping functions to re-scale the time axis on the raw data; it does *not* directly rely on the response template, 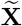. Thus, one can visualize the aligned activity of an individual neuron in a held out manner—the model is fit to all trials and all other neurons, and the warping functions are applied to the held out cell. This can then be repeated for each neuron in the full population. All spike raster plots in the main paper were produced using this procedure.

While these two approaches do not supplant the need for careful cross-validation, they can provide a quick validation for visualizations and presented results.

### Synthetic data examples

In **Figure 1** data from a single neuron was simulated as a difference of two exponential curves. The activity at *T* = 100 equally spaced time points between [−8, +8] was given by:

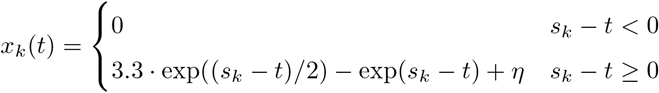

Where *s*_*k*_ was a random shift parameter drawn uniformly on the interval [−5.5, 3.5), and *η* was randomly drawn zero-mean gaussian noise with a standard deviation of 0.15. Unregularized shift-only, linear, and piecewise linear (with 1 knot) models were fit to *K* = 100 simulated trials. DTW-Barycenter Averaging (DBA; Petitjean et al. 2011) was fit to the same data using (Tavenard 2017).

In **Figure 2** we simulated random warping warping functions following the procedure listed in **Equation ??**, with *Q* = .12. The firing rate template of each neuron was given by a smoothed, sparse sequence of heavy-tailed random variables:

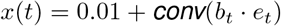

where *e*_*t*_ were randomly drawn from an exponential distribution (with scale parameter equal to one) and *b*_*t*_ were binary random variables drawn from a Bernoulli distribution (with probability of 0.92 that *b*_*t*_ = 0). The conv(·) procedure denotes convolution with a Gaussian smoothing kernel with a standard deviation of 2. Truncated Poisson random variables were then drawn in each timebin; any bins with more than two spikes were truncated to one spike.

## Experimental Methods

### Mouse Olfactory Task

All procedures were approved by the Institutional Animal Care and Use Committee of New York University Langone Medical Center. We analyzed data that was collected as part of a previously published study (Wilson et al. 2017).

Data presented from the mouse olfactory bulb were collected from a single recording session using an awake male C57B/6 mouse. Subject was implanted with a RIVETS headbar for head-fixation 7 days prior to the experiment (see description in Arneodo et al. 2018). Subjects were water deprived prior to the experiment and were administered water during random odor presentations to acclimate animals to the experimental apparatus.

On the day of experiment, the subject was anesthetized using isoflurane and a ~0.3 mm craniotomy was preformed to gain access to the dorsal olfactory bulb. NeuroNexus A2×32 probes were inserted approximately 500 *μ*m into the dorsal bulb to record from the mitral-tufted cell layer. After probe insertion, the subject was allowed to recover from anesthesia for 30 minutes prior to recording. Electrophysiological and respiration signals were recorded using the HHMI Janelia Whisper recording system at 25000 Hz. Respiration (sniff) was monitored non-invasively using a pressure sensor sampling from the airflow in front of the nose. Action potentials from the recording were identified and classified into units offline using Spyking Circus template-matching software (Yger et al. 2018).

Two odors at 3 concentrations were presented in randomly interleaved trials. Subjects were passively sampling odor during trials. Concentrations covered a range of 2 orders of magnitude of molarity in carrier air. Odorants were diluted in mineral oil, stored in amber volatile organic analysis vials, and delivered via a 8-odor olfactometer. Odorant concentrations were controlled using a combination of gas- and liquid-phase dilution. We restricted our analysis to subsets of trials with odorant concentrations of 10^−2^ M—the highest concentration analyzed in (Wilson et al. 2017).

### Primate Motor Task

All procedures and experiments were reviewed and approved by the Stanford University Institutional Animal Care and Use Committee. Two male rhesus macaque monkeys (*Macacca mulatta*), denoted monkey J and monkey U, were used in this study. The monkeys were 13 (J) and 7 (U) years old and weighed 16 kg (J) and 13 kg (U) at the time of these experiments.

Monkeys performed a standard center-out delayed reach task described previously in (Gilja et al. 2012; Ames et al. 2014). Targets were presented at (40, 80, 120) cm and 90 cm from the central starting location for monkeys J and U respectively. For monkey J, delay periods were evenly distributed between 300 and 700 ms (monkey J) with ~4.5% non-delay trials randomly interleaved. For monkey U, delays were randomly distributed between 350 and 600 ms on 87% trials, between 5 and 350 ms on 10% of trials, with 3% non-delay trials. Monkeys received a liquid reward upon touching and holding the cursor on the target. Movement during the delay period caused a trial failure and provided a brief automated time out (~1 s). Non-delay trials were not analyzed.

For monkey J, the virtual cursor and targets were presented in a three-dimensional environment (MusculoSkeletal Modeling Software, Medical Device Development Facility, University of Southern California). Hand-position data were measured at 60 Hz with an infrared reflective bead–tracking system (Polaris, Northern Digital). Behavioral control and neural decode were run on separate PCs using the Simulink/xPC platform (Mathworks) with communication latencies of less than 3ms. This system enabled millisecond timing precision for all computations. Visual presentation was provided via two LCD monitors in a Wheatstone stereotax configuration, with refresh rates at 120 Hz, yielding frame updates of 7 ± 4 ms. Two mirrors visually fused the displays into a single three-dimensional percept for the user, as described previously in (Gilja et al. 2012).

For monkey U, the virtual cursor and targets were presented on a standard 2D display. The monkey controlled the position of an onscreen cursor using a haptic manipulandum which applied no additional forces applied to the arm and was only used for positional cursor control. The haptic device was constrained to move within a 2D vertical workspace and cursor position tracks hand position 1:1 without perceptible lag.

Neural recordings were obtained via implanted 96-electrode Utah Microelectrode arrays (Blackrock Microsystems) using standard neurosurgical techniques. Two arrays were implanted in the left hemisphere of Monkey J, one in dorsal premotor cortex (PMd) and one in primary motor cortex (M1). Three arrays were implanted in the left hemisphere of Monkey U, one in PMd, one in medial M1, and one in lateral M1. For both monkeys, implantation location was estimated visually from local anatomical landmarks.

Neural data were band-pass filtered between 250-7500 Hz, and processed to obtain multiunit ‘threshold crossings’ spikes, defined as any time the signal crosses −3.5 times RMS voltage. We did not perform spike sorting, and instead grouped together the multiple neurons present on each electrode. As such, we anticipate that these population recordings contain both single and multiunit activity.

For **Figure 4** and **Figure 4, Supplement 1**(Monkey J), each trial was defined as the 1200 ms following the reach target onset. Spike times were binned in 5 ms increments. For **Figure 5**(Monkey U) and **Figure 5, Supplement 1** (Monkey J), we aligned spike times to the go cue instead of target onset. To highlight oscillatory spiking activity, we defined each trial as the period occurring 400 ms prior to go cue and 100 ms after go cue. Spike times were binned in 2.5 ms increments for Monkey J and 5 ms increments for Monkey U; similar results were found for smaller bin sizes, and stronger smoothness regularization. In **Figure 5F**(Monkey U), we extended each trial duration to *±*400 ms around the go cue, but otherwise kept the same parameters.

Tuning the regularization strength of on template smoothness (*λ*) and warp magnitude (*μ*) was important to uncover the oscillations in premotor cortex. We used the cross-validation procedure described above to determine roughly appropriate values for these parameters; we increased the regularization strength further for the purposes of visualization and to be confident that are results were not due to overfitting.

### Rat Motor Task

All procedures and experiments were reviewed and approved by the Harvard Institutional Animal Care and Use Committee. We analyzed data that was collected as part of a previously published study (Dhawale et al. 2017), which describes all experimental procedures and data collection protocols in greater detail.

Experimental subjects were female Long Evans rats, 3-8 months old at the start of the experiment (Charles River). Extracellular recordings were obtained from 16 chronically implanted tetrodes in the motor cortex. Signals were amplified and digitized on a customized head-stage, and sampled at 30 kHz. The head stage was attached to a custom-designed tethering system that allowed the animal to move freely within its cage. Before implantation, an automated behavioral training framework (described in Poddar et al. 2013) was used to train the rats on a timed lever-pressing task (described in Kawai et al. 2015) until asymptotic performance was achieved.

The tetrode drive was then surgically implanted and targeted to motor cortex, through a 4-5 mm diameter craniotomy made 2 mm anterior and 3 mm lateral to bregma. The tetrode array was lowered to a target depth of 1.85 mm. At the end of the experiments, the position of the electrodes was verified by standard histological methods—brains were fixed via transcardial perfusion (4% paraformaldehyde in phosphate-buffered saline, Electron Microscopy Sciences) and the location of the electrodes was reconstructed by viewing mounted coronal sections (60 mm).

After 7 days of recovery post-surgery, training on the task resumed in the animal’s home cage. Neural and behavioral data was recorded continuously during this time (12-16 weeks) with only brief interruptions (median time of 0.2 hr). Spikes were sorted using Fast Automated Spike Tracker (FAST), a custom algorithm designed for parsing long-term continuous neural recordings (for details, see Dhawale et al. 2017). We examined *K* = 1265 trials, collected over a two day period.

Each trial was defined as the period starting 500 ms prior to the first lever press and 1500 ms after the first lever press. Spike times were binned in 10 ms increments for each unit. Raw spike counts were provided to the time warping algorithm; however, we observed similar results under various normalization schemes, such as soft-normalization (Churchland et al. 2012). All analyses of these data used a shift-only time warping model. The per-trial shift was constrained to be less than 10% of the total trial duration.

## Supplemental Figures

**Figure 3, Supplement 1.**
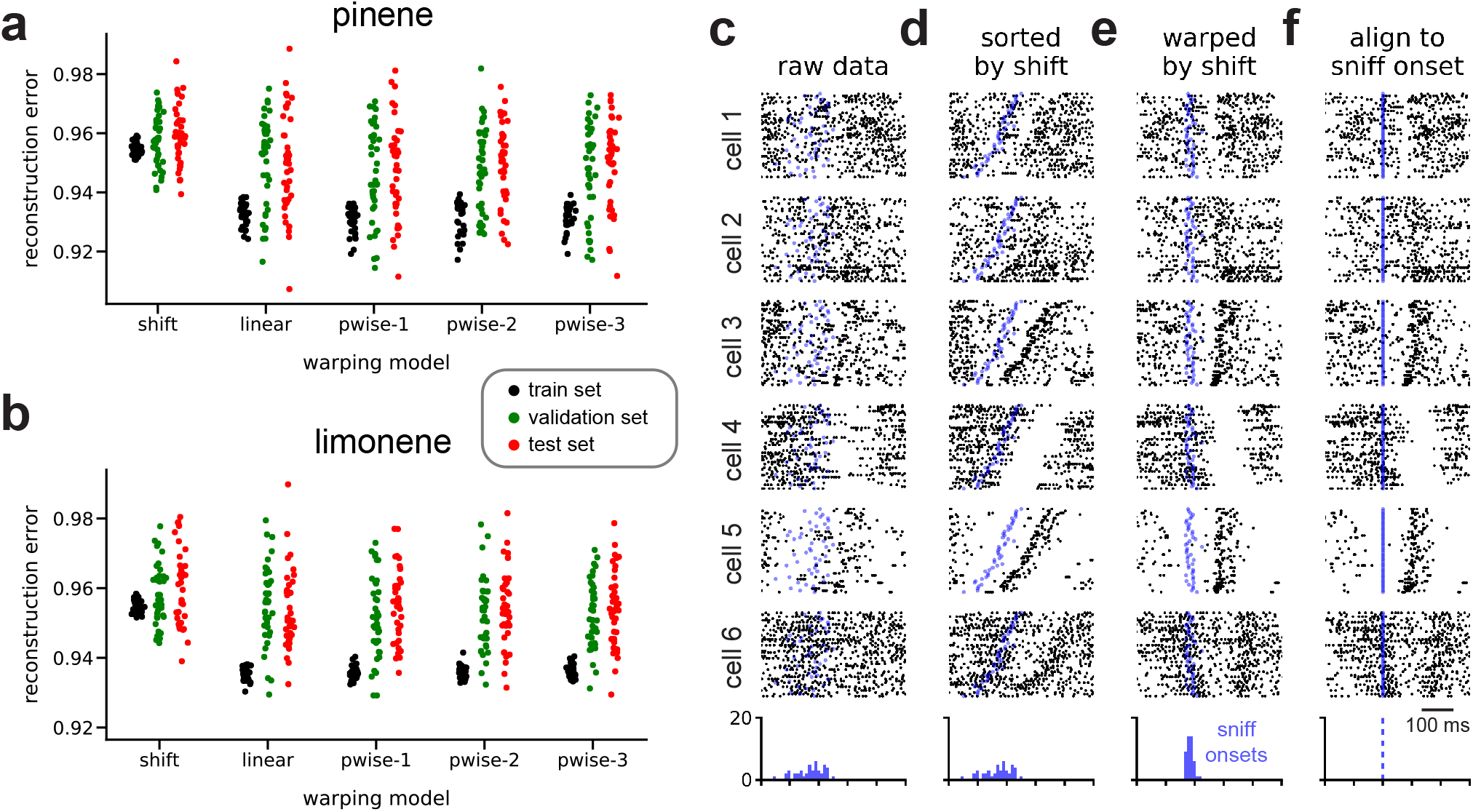
Nested cross-validation results and supplementary results on a different odorant. (A) Nested cross-validation results for data presented in Fig. 2 (pinene, 10^−2^ M). Vertical axis shows Euclidean norm of model residuals divided by norm of data for training and test sets. (B) Same as panel A, but computed on neural responses to a different odorant (limonene, 10^−2^ M). (C-F) Same as Fig. 2B-E in the main text, but on *K* = 45 presentations of limonene (10^−2^ M).

**Figure 4, Supplement 1.**
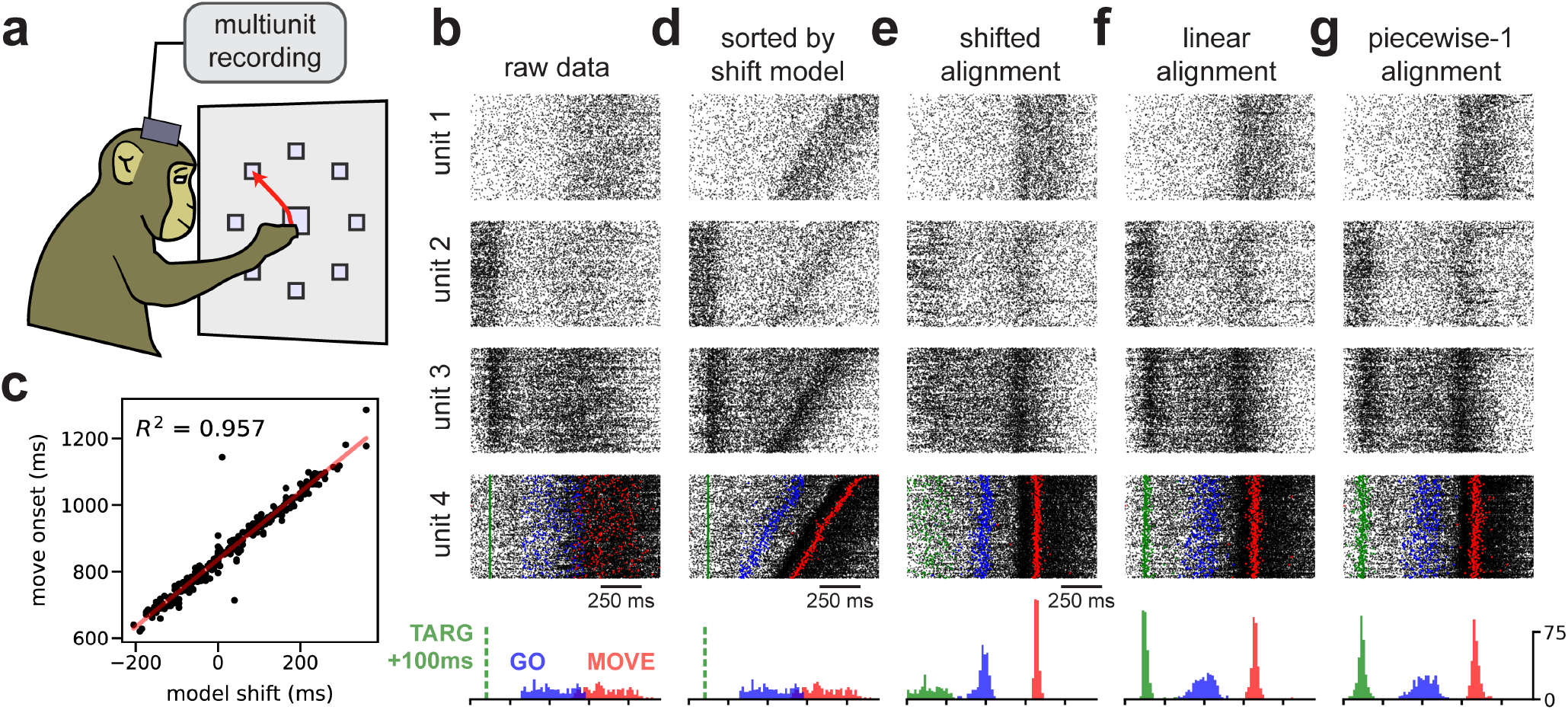
Replication of movement onset detection in primate reaching experiment. All figure panels are directly analogous to **Figure 4** in the main text and show comparable results. Instead of 90° reach trials (analyzed in the main text) only 135° reach trials were analyzed.

**Figure 4, Supplement 2.**
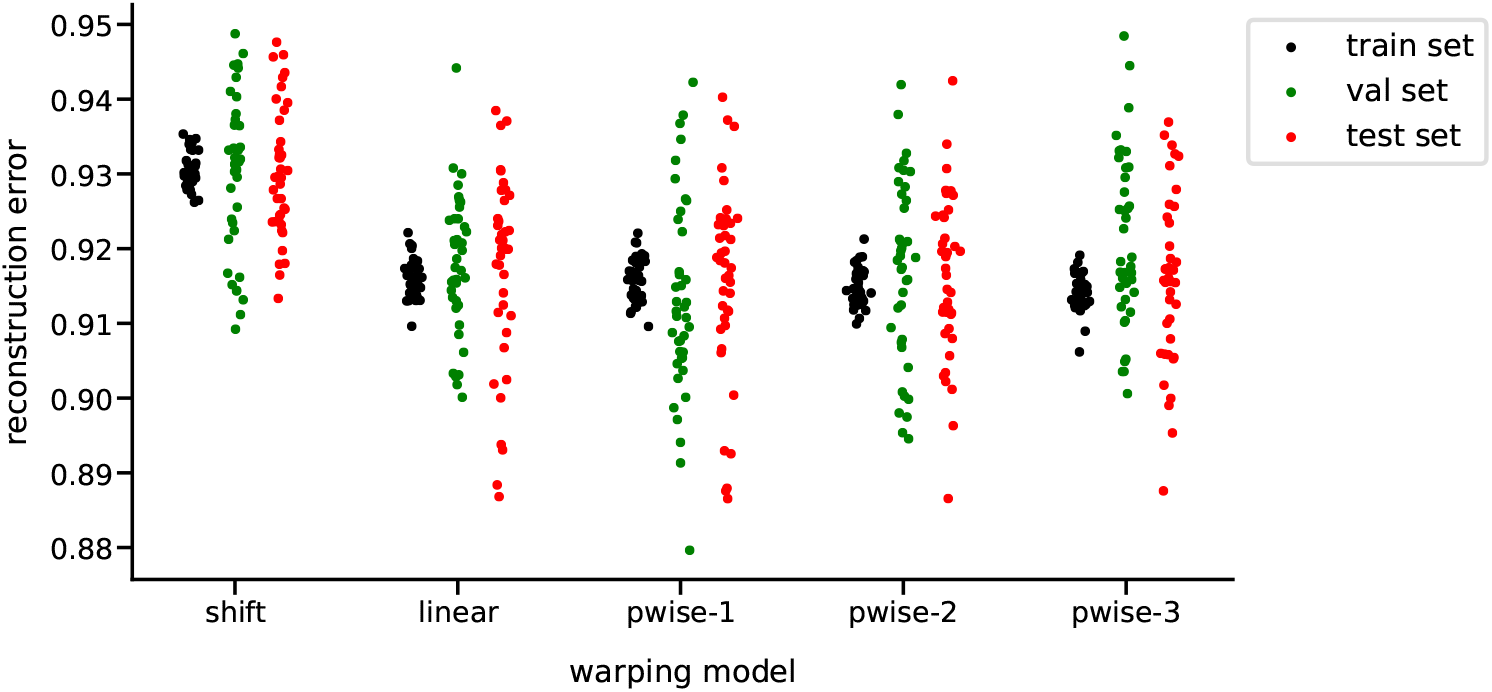
Nested cross-validation of primate reaching dynamics aligned to target onset.

**Figure 5, Supplement 1.**
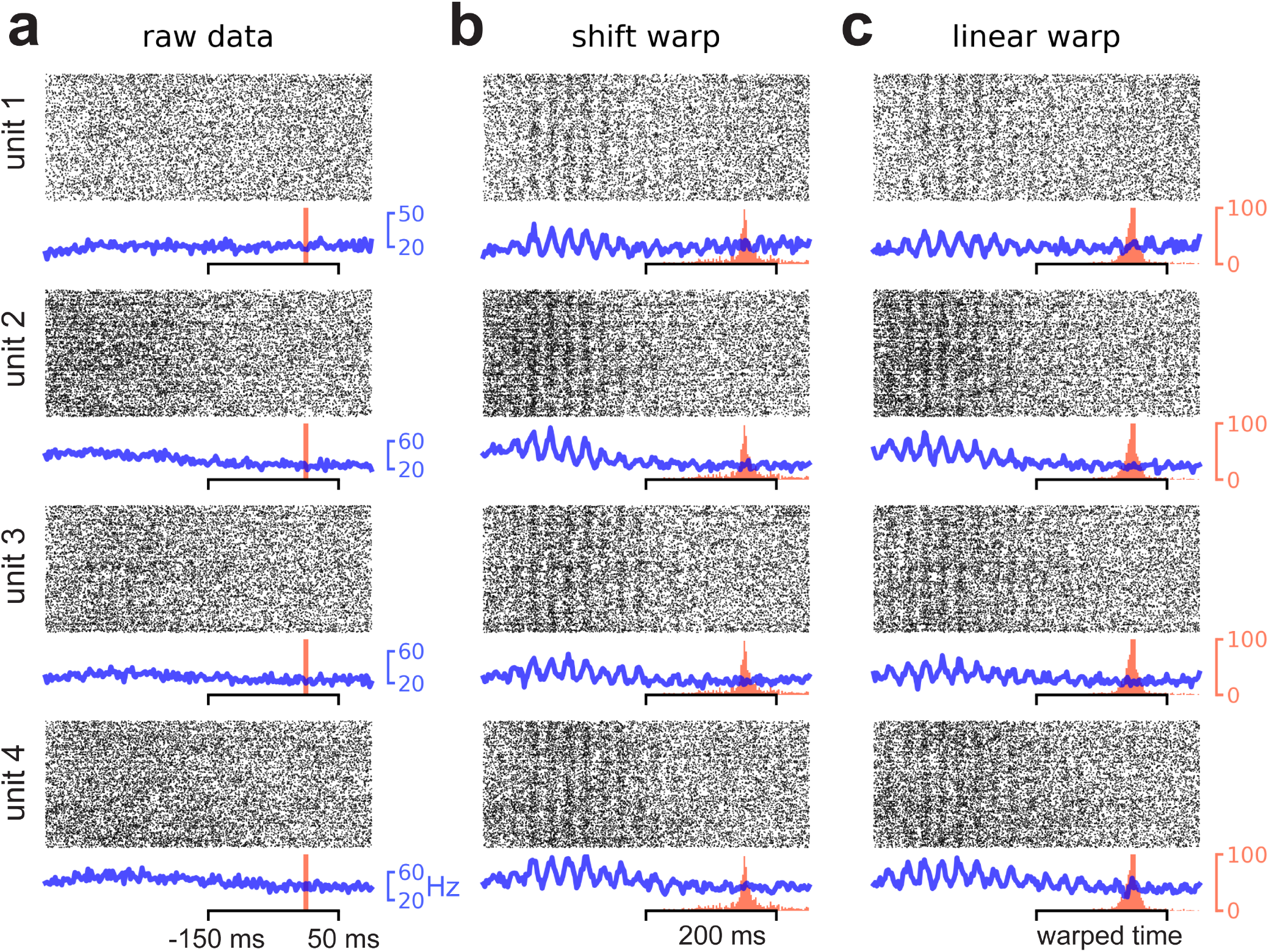
Oscillations in premotor cortex uncovered in data from a second nonhuman primate. (A) Trial-by-time raster plots (black) and trial-averaged estimates of firing rate (blue) for four example multiunits. Red vertical line denotes the time that the go cue was delivered. (B-C) Same as panel A, except after spike times aligned by shift-only time warping (B) or linear time warping (C). Red histogram shows the distribution of go cue times after after the time warping transformation was applied. All four multiunits were held out during model fitting—the warping functions were fit to the remaining *N* = 95 units and applied to the held out multiunit to generate the displayed raster plots.

**Figure 5, Supplement 2.**
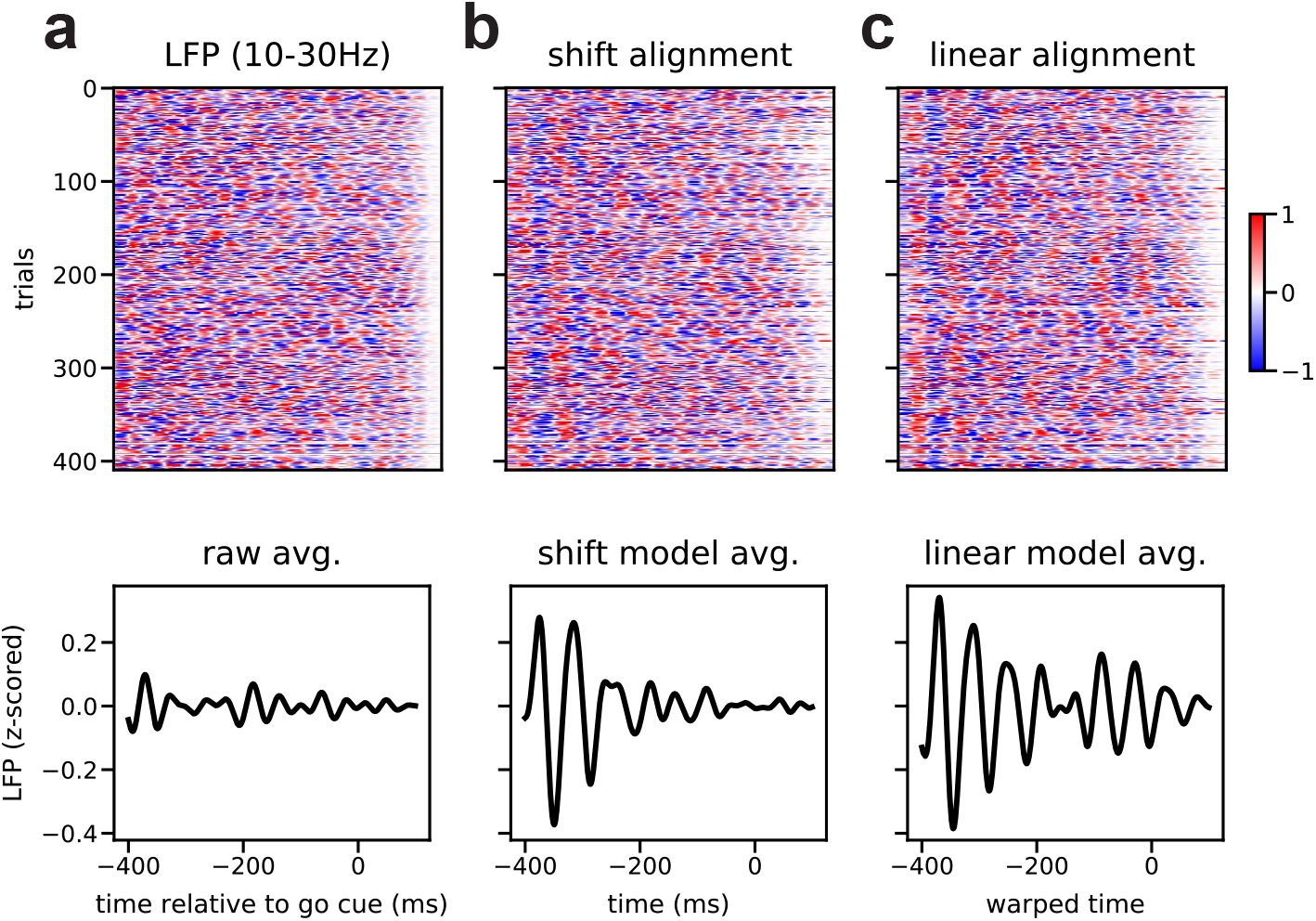
Spike time oscillations in primate premotor cortex align with LFP. (A) Top, LFP signal on all trials. The signal was obtained by averaging over all *N* = 96 electrodes, z-scoring the signal within each trial, and then bandpass filtering (10-30 Hz; fifth-order Butterworth digital filter). Bottom, average LFP signal across trials. (B-C) Same as panel A, except after applying temporal alignments from a shift-only warping model (B) and a linear warping model (C). In both cases, time warping uncovered strong oscillations at ~18 Hz—the same frequency of spike-level oscillations identified in Fig 4. Importantly, the warping models were fit *only* to binned spike times, demonstrating that the model generalized well to new data stream with fundamentally distinct features. This suggests that the spike-level oscillations described in Fig 4 are time-locked with LFP oscillations, in agreement with prior work (Murthy and Fetz 1992; Sanes and Donoghue 1993; Reimer and Hatsopoulos 2010; Pandarinath et al. 2018). All data were taken from the same animal subject shown in Fig 4.

**Figure 6, Supplement 1.**
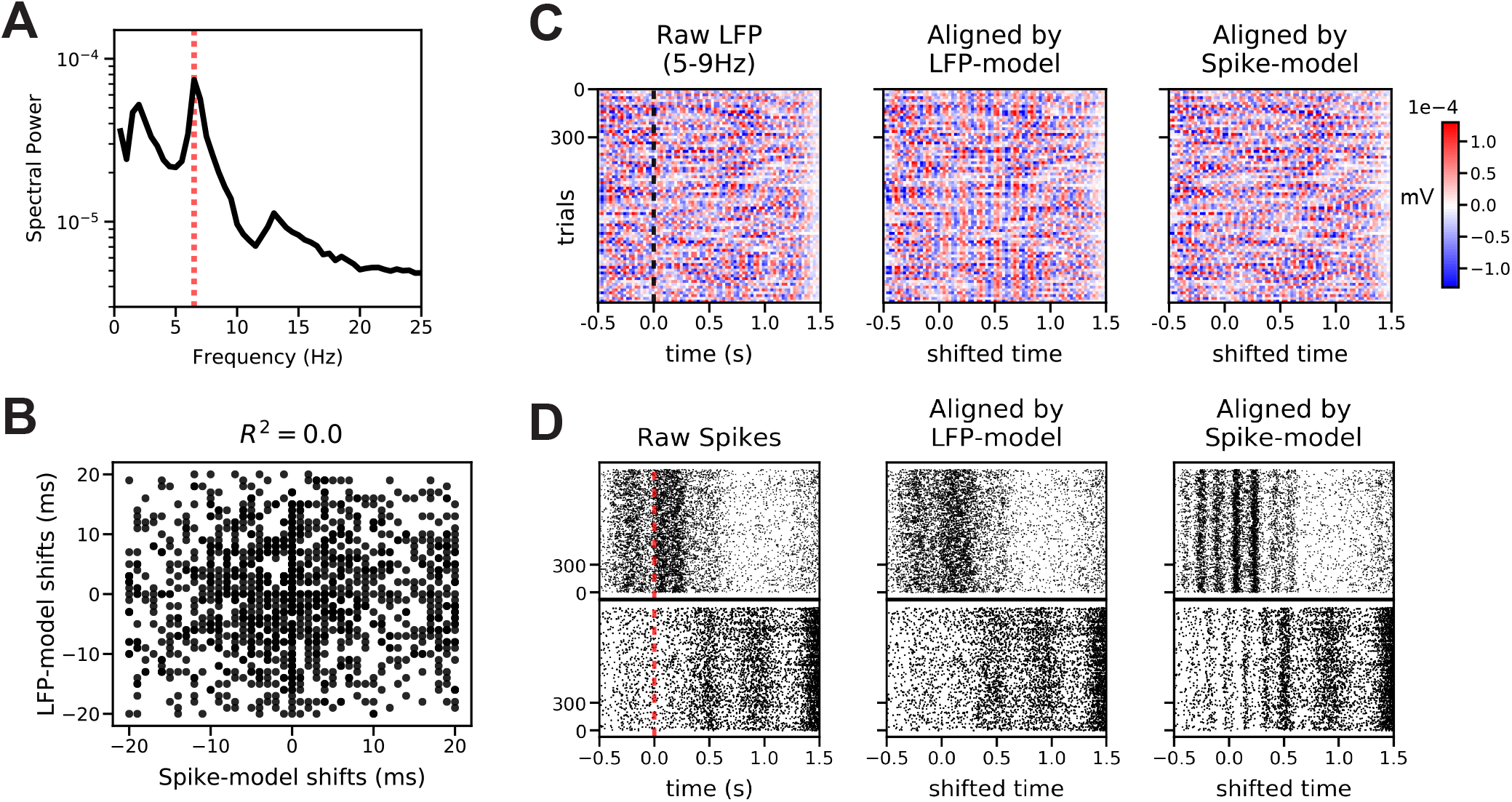
LFP does not correlate with spike-level oscillations in rat motor cortex. The LFP signal was highly correlated across all electrodes and thus averaged across electrodes before analysis. (A) Trial-averaged periodogram of the LFP signal. Dashed red line denotes 6.5Hz, illustrating a peak in the LFP spectrum that is similar to the frequency of spike-level oscillations in **Fig 6**. (B) One shift-only time warping model was fit to bandpassed-filtered LFP signals (LFP-model; fifth-order digital Butterworth, 5-9 Hz), and a second shift-only time warping model was fit to binned spike trains (Spike-model; same as **Fig 6**). The scatterplot demonstrates the per-trial shift parameters learned by these models were not correlated, suggesting that the spike-level oscillations are not phase-locked to LFP. (C) Bandpassed LFP as raw data (left; dashed line denotes first lever press), and same data aligned by LFP-model (middle) and Spike-model (right) time warping models. LFP is not reliably aligned by the time warping model fit to spiking data. (D) Raster plots from two example neurons (top and bottom rows), showing raw spike times (left; dashed line denotes first lever press), aligned by LFP-model (middle), and aligned by Spike-model (right). Spike-level oscillations are not revealed by the time warping model fit to LFP.

**Figure 6, Supplement 2.**
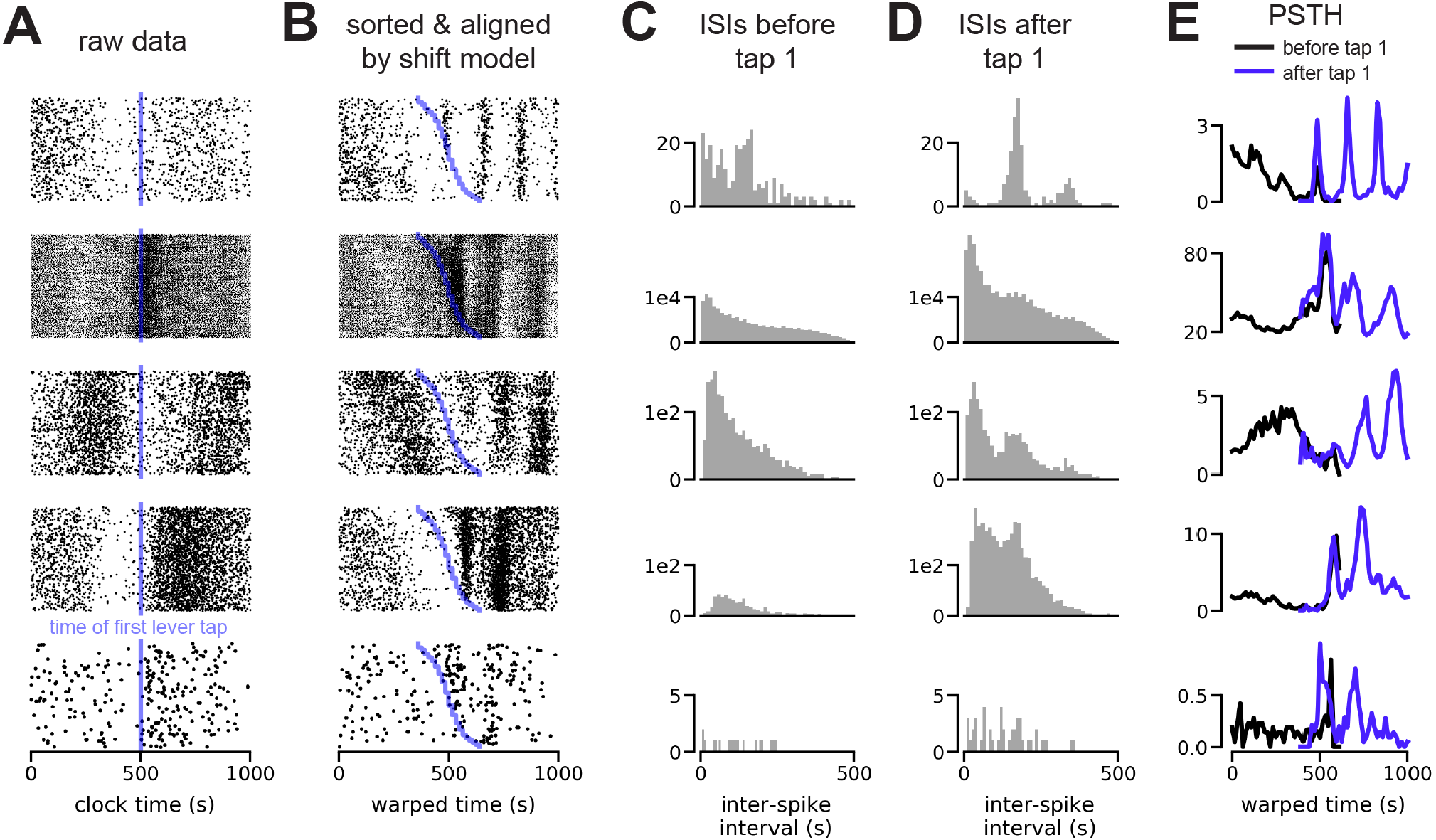
Five representative isolated units exhibiting stronger spike time oscillations following the first lever press. (A) Raw spiking activity in a 1 second window around the first lever press. Vertical blue line denotes the time of the first lever press (manual alignment point). (B) Model-aligned spiking activity by shift-only warping, with trials sorted by the direction and magnitude of the learned shift. Blue line denotes the time of the first lever press on each trial. (C) Inter-spike interval (ISI) distributions during the 500 ms preceding the first lever press. (D) ISI distributions during the 500 ms following the first lever press. Note increased peak around ~150 ms, corresponding to increased oscillations at ~7 Hz. (E) Trial-averaged PSTHs for model-aligned spike times. Black lines denote PSTHs computed from spikes preceding the first lever press, while blue lines denote PSTHs computed from spikes following the first lever press. Note increased oscillatory dynamics following the lever press.

